# pH adjustment increases biofuel production from inhibitory switchgrass hydrolysates

**DOI:** 10.1101/2025.01.10.632484

**Authors:** Lillian M. Barten, Johnathan G. Crandall, Dan Xie, Jose Serate, Evan Handowski, Annie Jen, Katherine A. Overmyer, Joshua J. Coon, Chris Todd Hittinger, Robert Landick, Yaoping Zhang, Trey K. Sato

**Author notes:** Corresponding Author, 1552 University Ave., Madison, WI 53726. These authors contributed equally (Lillian M. Barten, Johnathan G. Crandall). Deceased (Yaoping Zhang).

## Abstract

Biofuels derived from renewable and sustainable lignocellulosic biomass, such as switchgrass, offer a promising means to limit greenhouse gas emissions. However, switchgrass grown under drought conditions contains high levels of chemical compounds that inhibit microbial conversion to biofuels. Fermentation of drought switchgrass hydrolysates by engineered *Saccharomyces cerevisiae* and *Zymomonas mobilis* generates less ethanol than fermentation of hydrolyzed switchgrass from an average rainfall year. Here, we demonstrate that this inhibitory effect can be alleviated by altering the pH of drought-switchgrass hydrolysates made from two different pretreatment methods: Ammonia Fiber Expansion (AFEX) and Soaking in Aqueous Ammonia (SAA). Fermentation rates and biofuel production from AFEX- and SAA-pretreated switchgrass hydrolysates from normal and drought years were higher at pH 5.8 than at pH 5.0 for both S*accharomyces cerevisiae* and *Zymomonas mobilis*. Additionally, SAA pretreatment of drought switchgrass enabled increased fermentation rates and titers compared to AFEX pretreatment. Using a synthetic mimic of switchgrass hydrolysate, we identified relief from pH-dependent inhibition by lignocellulose-derived inhibitors as the cause of increased biofuel production above a pH of 5.0. These results demonstrate that SAA pretreatment and pH adjustment can significantly improve fermentation and biofuel production from switchgrass hydrolysates and especially from drought-switchgrass hydrolysates by industrial microorganisms.

## 1. Introduction

Lignocellulosic-derived biofuels are being investigated as a sustainable and renewable solution for combatting climate change and reducing greenhouse gas emissions. Although electrification has provided an alternative carbon-neutral energy source to replace traditional liquid automobile fuels, electrification is a less viable option for the aviation industry. Sustainable aviation fuels (SAF) generated from alcohol-to-jet (ATJ) processes are one of many proposed approaches to decarbonize the aviation sector (Geleynse et al., 2018; Holladay et al., 2020). These alcohols include ethanol, which can be catalytically converted into longer chain ethers, olefins, and paraffins (Xie et al., 2024), as well as isobutanol, which can be converted to branched alkenes and alkanes, and cyclic molecules (Geleynse et al., 2018; Holladay et al., 2020). Both ethanol and isobutanol can be produced through microbial fermentation of renewable lignocellulosic feedstocks. Potential plant feedstocks include switchgrass, a drought- and heat-tolerant grass species with resistance to insects and disease that can grow on agricultural land unsuitable for food production (Keshwani & Cheng, 2009; Rinehart, 2006). Moreover, switchgrass creates over 500% more energy than is used to produce it (Schmer et al., 2008), making switchgrass a promising feedstock for ATJ biofuels.

Like all living organisms, switchgrass responds to both abiotic and biotic stressors, which induce physiological responses that affect their biomass compositions. These stressors, including extreme temperatures, floods, and drought can have downstream consequences on biofuel production by inhibiting conversion microbes such as the yeast *Saccharomyces cerevisiae* and bacteria *Zymomonas mobilis*. In 2012, a severe drought in the Midwest (Mallya et al., 2013) impacted the composition of switchgrass grown in Wisconsin, causing the biomass to accumulate elevated levels of osmoprotectant sugars (Ong et al., 2016) and saponins (Chipkar et al., 2022). When deconstructed with Ammonia Fiber Expansion (AFEX) pretreatment and enzymatic hydrolysis, the 2012 switchgrass hydrolysate was highly inhibitory to fermentation by *S. cerevisiae* but not by *Z. mobilis* (Chandrasekar et al., 2021; Chipkar et al., 2022; Ong et al., 2016). Rates of yeast cell growth, ethanol production, and sugar consumption were significantly lower in hydrolysates from 2012 switchgrass than in fermentations of deconstructed switchgrass grown in 2010 and 2016, which had higher seasonal rainfall. Chemical analysis determined that hydrolysates from deconstructed 2012 switchgrass contained higher levels of pyrazines and imidazoles, which can have inhibitory effects on yeast (Ong et al., 2016). Switchgrass biomass from 2012 also contained higher amounts of multiple saponins—plant-derived, bioactive secondary metabolites—than the 2010 and 2016 feedstocks (Chipkar et al., 2022). This result suggests there are multiple compounds causing these inhibitory effects in yeast fermentation, potentially working together to inhibit fermentation in the 2012 drought–switchgrass hydrolysate.

In addition to nitrogenous compounds and saponins, deconstructed switchgrass hydrolysates contain numerous other chemical compounds that have varying effects on microbial fermentation and biofuel production. Some of these inhibitory compounds are directly produced by the plants whereas others, such as phenolic compounds, are generated during the pretreatment process (van der Pol et al., 2014). The mechanisms by which all these compounds inhibit biofuel-producing yeast and bacteria are only somewhat understood. For example, saponins are predicted to have pore-forming effects on microbial cell membranes (Mugford & Osbourn, 2013), perhaps due to interactions with ergosterol in the cell membrane (Alcázar et al., 2017; Sokolov et al., 2022). Some known inhibitory compounds, such as weak acids (acetic acid), furans (furfural), and phenolic acids (ferulic and coumaric acids), can damage membranes, decrease cellular pH, and induce depletion of cellular ATP (Piotrowski et al., 2014). Increasing the pH of some lignocellulosic hydrolysates with the addition of potassium can alleviate inhibitory effects of furfural and other toxins, resulting in increased ethanol production (Lam et al., 2021).

Here, we explored whether the inhibitory effect of 2012 switchgrass was specific to the AFEX pretreatment process, and whether this effect could be alleviated by pH adjustment to the pretreated hydrolysate. We hypothesized that increasing the pH of hydrolysate to at or near the pK_a_ of some key acidic inhibitors would have beneficial effects on microbial fermentation. Switchgrass from drought (2012) and normal precipitation years (2010 and 2016) were pretreated using both AFEX and Soaking in Aqueous Ammonia (SAA) methods followed by enzymatic hydrolysis. The pH of hydrolysates produced using both pretreatment methods was adjusted to 5.0 and 5.8, and these feedstocks were fermented by both *S. cerevisiae* and *Z. mobilis*. pH adjustment enabled substantially higher biofuel production rates from both hydrolysates, as well as from two formulations of synthetic switchgrass hydrolysate.

## 2. Materials and Methods

### 2.1 Feedstocks

Switchgrass grown in 2010, 2012, and 2016 were harvested, dried, and milled to 5 mm particle size as described previously (Ong et al., 2016; Serate et al., 2015; Zhang et al., 2020). Glucan contents (%) for the 2010, 2012, and 2016 switchgrass were previously determined to be 34.8%, 30.5%, and 34.5%, respectively (Ong et al., 2016; Zhang et al., 2020).

### 2.2 Biomass pretreatment

AFEX pretreatment of switchgrass were described previously for the 2010 and 2012 harvests (Serate et al., 2015) and for the 2016 harvest (Zhang et al., 2020). The Soaking in Aqueous Ammonia (SAA) pretreatment method was developed based on a previously published patent (Dunson Jr. et al., 2007). 15 g of milled switchgrass was weighed out and transferred into a 120 mL glass pressure tubes (Chemglass, Cat# CG 1880-06). 60 mL of 18% diluted ammonium hydroxide (Fisher Chemical, Cat# A669-212) was slowly added to each tube, with mild agitation if needed, and the tubes were then sealed tightly. Sealed tubes were left upside down for 1-2 minutes and then mixed by inversion to ensure saturation of the biomass with ammonium hydroxide. Tubes were then placed in a rotator set at 19 RPM within a hybridization oven (ProBlot) set at 75°C. After 1.5 hr incubation, tubes were removed and mixed, then returned to the 75°C oven for an additional 2 hr of incubation (3.5 hr total incubation time at 75°C). After incubation at 75°C for 3.5 hr, tubes were placed inside of a fume hood for 30 minutes too cool. After the cooling period the caps were carefully unscrewed to release the pressure.

Drying of SAA-pretreated samples was handled in two stages. First, pressurized drying with compressed air was performed with a rotary meter. The clear silicone tubing that connects house air and the rotary meter is set to ∼20 L/min. A metal spatula was used to make a gap in the center of the biomass to the bottom of the glass tube for the silicone tubing. The end of the tubing was then placed ∼0.25 inches from the bottom of the glass tube and air was blown into the sample (“pressurized drying”). After 48 hr of pressurized drying, the pretreated biomass was transferred from the glass pressure tube to weigh boats and further dried in a fume hood for 5 days. Dried SAA-pretreated biomass was then transferred to a glass bottle for storage until enzymatic hydrolysis. A minimum of 0.5 g of dried SAA-pretreated biomass was used for moisture content analysis (Ohaus MB35 Moisture Analyzer).

### 2.3 Enzymatic hydrolysis of pretreated switchgrass

AFEX- and SAA-pretreated switchgrass samples were loaded into 85mL Oak Ridge tubes (Thermo Scientific, Nalgene, Cat# 3118-0085) for 7% glucan loading in 50 mL total volume. Pretreated biomass was brought to 32.7 mL with double-deionized water (ddH_2_O) and mixed with 5 mL 1M potassium phosphate buffer, and then autoclaved. After the biomass was cooled, 150 µL of HCl (∼37–38% w/v) was added and mixed, and then 5.8 mL of a sterile-filtered enzyme mixture (1 mL cellulase cocktail, Novozymes NS 22257; 0.17 mL xylanase cocktail, Novozymes NS 22244; 4.7 mL ddH_2_O) was added. Sealed tubes were mixed and incubated on a rotator at 50°C. After 2 days of hydrolysis, a second loading of 5.8 mL enzyme mixture was added to the SAA-pretreated biomass, whereas the AFEX-pretreated material received 5.8 mL of ddH_2_O. After 5 additional days of hydrolysis at 50°C, the hydrolysate was centrifuged at 15,000 x *g*, pH adjusted to 5.0 with NaOH or HCl, and then sterile filtered with a 0.2-micron filter unit (Nalgene). When needed, half of the volumes of each hydrolysate at pH 5.0 were further adjusted to pH 5.8 with NaOH.

### 2.4 Analysis of hydrolysate chemical compositions

Detection and quantification of chemical compounds in hydrolysates were determined by liquid chromatography-mass spectrometry/mass spectrometry (LC-MS/MS) as described previously (Zhang et al., 2020). In brief, hydrolysate samples were diluted ten□fold with chilled μM vanillin-^13^C_6_ in water, mixed well by pipetting up and down, centrifuged at 14,000 x g for 2 minutes at 4°C, and then the supernatant was transferred to an amber glass autosampler vial with glass vial insert for LC□MS/MS analysis. Standard calibration curves for quantitation of 84 chemical compounds were generated with twelve total standard concentration levels ranging from 0.024 μM to 50 μM. Each standard level was transferred to a low volume amber autosampler vial with insert and then placed into an autosampler at 4°C for injection.

LC□MS/MS analyses were performed with a 5 μL injection volume by Vanquish Split Sampler HT autosampler (Thermo Scientific). Sample components were separated using an ACQUITY UPLC HSS T3 reversed phase column (2.1 x 150 mm, 1.8 μm particle diameter, Waters Corporation) held at 40°C, and a Vanquish Binary pump (150 μL/min flow rate; Thermo Scientific). Mobile phase A consisted of 0.1% formic acid in water. Mobile phase B consisted of 100% acetonitrile. Mobile phase B was increased from 7.5% to 30% over 20 min. Mobile phase B was further increased to 100% over 14.5 min and then held at 100% for 1.5 min. The column was re□equilibrated with mobile phase B at 7.5% for 2 minutes before the next injection.

The LC system was coupled to a Q Exactive Orbitrap mass spectrometer through a heated electrospray ionization (HESI II) source (Thermo Scientific). Source conditions were as follows: HESI II and capillary temperature at 275°C, sheath gas flow rate at 30 units, auxiliary gas flow rate at 6 units, sweep gas flow rate at 0 units, spray voltage at |4.0k| for positive mode and |4.5k| for negative mode, and Slens RF at 60.0 units. To promote fragmentation, in□source collision□induced dissociation (CID) energy was set to 25.0 eV. The MS was operated in a polarity switching mode acquiring positive and negative full MS and targeted MS2 spectra (top 2) within the same injection. Acquisition parameters for full MS scans in both modes were 35,000 resolution, 1 x 105 automatic gain control (AGC) target, 50 ms ion accumulation time (max IT), and 50□360 m/z scan range. Targeted MS-MS scans in both modes were then performed at 17,500 resolution, 1 x 105 AGC target, 100 ms max IT, 1.0 m/z isolation window, stepped normalization collision energy (NCE) at 20, 30, 40, and a 5.0 s dynamic exclusion. The retention time window for each compound was optimized from prior experiments. The resulting LC□MS/MS data was manually processed using a custom TraceFinder 4.1 (Thermo Scientific) method using a mass precision of 4 ppm and mass tolerance of 10 ppm. The prepared standard solution was used to locate and identify appropriate peaks for peak area analysis.

### 2.5 Media and microbial strains

The xylose-fermenting *Saccharomyces cerevisiae* strain GLBRCY1455 was derived from the strain GLBRCY1327 (Lee et al., 2021) with an additional deletion mutation in *FLO8* to limit flocculation. *FLO8* was deleted by homologous recombination and marker rescuing with the *hphMX* cassette (Güldener et al., 1996). The generation of the isobutanol/ethanol co-producing hybrid *S. cerevisiae* strain yHRW253 was described previously (Pastore de Lima et al., 2023). Both *S. cerevisiae* strains were cultured in YPD medium (10 g/L yeast extract, 20 g/L peptone, and 20 g/L dextrose). Xylose-fermenting *Zymomonas mobilis* 2032 (Yang et al., 2018) obtained from the American Type Culture Collection, PTA-6977) was cultured in *Zymomonas* Rich Defined Medium (ZRDM; (Enright et al., 2023)) with tetracycline (Tc) and chloramphenicol (Cm). Synthetic hydrolysate versions 4.0 and 4.1 (SynHv4.0 and SynHv4.1) mimic the composition of 7% glucan loading AFEX-pretreated switchgrass as previously determined (Serate et al., 2015; Zhang et al., 2020). SynHv4.1 contains specific lignocellulose-derived inhibitors (*e.g.*, ferulic and p-coumaric acids), whereas SynHv4.0 lacks these inhibitors. Chemical composition of SynHv4.0 can be found in Supplemental File S1. The composition of SynHv4.1 is the same as SynHv4.0 but contains additional lignocellulose-derived inhibitory compounds (Supplemental Table S2). Tween-80 and ergosterol were made by dissolving ergosterol (E6510-10G, Sigma) and Polysorbate 80 (Tween 80, 59924-1KG-F, Fluka) in 100% ethanol. The pH of all media and hydrolysates were adjusted with HCl and NaOH to pH values of 3-7 as needed.

### 2.6 Hydrolysate fermentations

Fermentations of hydrolysates by yeast and bacteria were performed as described in (Chandrasekar et al., 2021) with modifications. *S. cerevisiae* and *Z. mobilis* cultures were grown overnight in YPD and ZRDM + Tc + Cm respectively. The next day, cultures were diluted to an optical density at 600 nm wavelength (OD_600_) = 0.3 with fresh media and grown for approximately 3-5 hours until cultures reach log phase at OD_600_ = 0.6-0.8. Cultures were then harvested and used to inoculate 4 mL of hydrolysate in sterile serum bottles (Fisher: 06406H) at an OD_600_ = 0.2. Inoculated bottles were sealed with 20 mm butyl rubber stoppers (ChemglassLife Sciences) and rendered anaerobic by three alternating vacuum and N_2_ gas-sparging cycles with syringe needles and tubing. Anaerobic cultures were connected to the Challenge Technology (Springdale, AR, USA) AER-800 respirometer system by piercing the rubber stoppers with syringe needles and tubing. The respirometer measures and recovers the volume of CO_2_ gas produced over time. Inoculated cultures connected to the respirometer were shaken at 150 RPM at 30°C for 48-72 hours. At the end of the experiment, cultures were harvested to measure the final cell density and centrifuged to collect the supernatant for metabolomic analysis. Supernatant glucose, xylose, and ethanol concentrations were determined by high-performance liquid chromatography (HPLC) and refractive-index detection (RID) as previously described (Schwalbach et al., 2012). Isobutanol concentrations were determined by gas chromatography as previously described (Gambacorta et al., 2022). Visualization and statistical analyses of growth metrics, metabolites, and hydrolysate composition were performed in R v4.4.1.

## 3. Results

### 3.1 Soaking in Aqueous Ammonia and AFEX pretreatments yield switchgrass hydrolysates with similar sugar titers but different molecular compositions

We first sought to determine if the inhibitory effect of the 2012 switchgrass hydrolysate would persist for a pretreatment different from AFEX, which is known to generate higher levels of phenolic amides (Keating et al., 2014; Zhang et al., 2020). Soaking in Aqueous Ammonia (SAA) pretreatment (Zhao et al., 2020) has been used to effectively deconstruct corn stover (Kim & Lee, 2007), rice straw (Ko et al., 2009), and switchgrass (Pryor et al., 2011), and it has been patented for industrial processes (Dunson Jr. et al., 2007). We customized this method to pretreat small amounts (10–15 g) of biomass in batches of 12–24 samples at a time. For this SAA-deconstruction pipeline (**Fig. 1A**), milled switchgrass samples harvested in 2010, 2012, and 2016 were incubated with mixing in 18% (wt/vol) NH_4_OH for 3.5 hr at 75°C in 120 mL glass pressure tubes, then cooled and dried with compressed air in a fume hood (see Materials and Methods section). Dried SAA-pretreated switchgrass samples at 7%-glucan loading were enzymatically hydrolyzed to produce 20–27 mL of filtered hydrolysate. To facilitate direct comparisons, AFEX-pretreated switchgrass samples from the same harvest years were hydrolyzed in parallel. Our SAA pretreatment conditions differred from previous AFEX pretreatment conditions used to deconstruct 2010, 2012, and 2016 switchgrass in a number of ways. These included different reactor vessel scales, ammonia–ammonium loadings, temperatures, pressures, and pretreatment times (**Table 1**). We expected that the differences in pretreatment conditions would result in hydrolysates with distinct compositions.

**Fig. 1.**
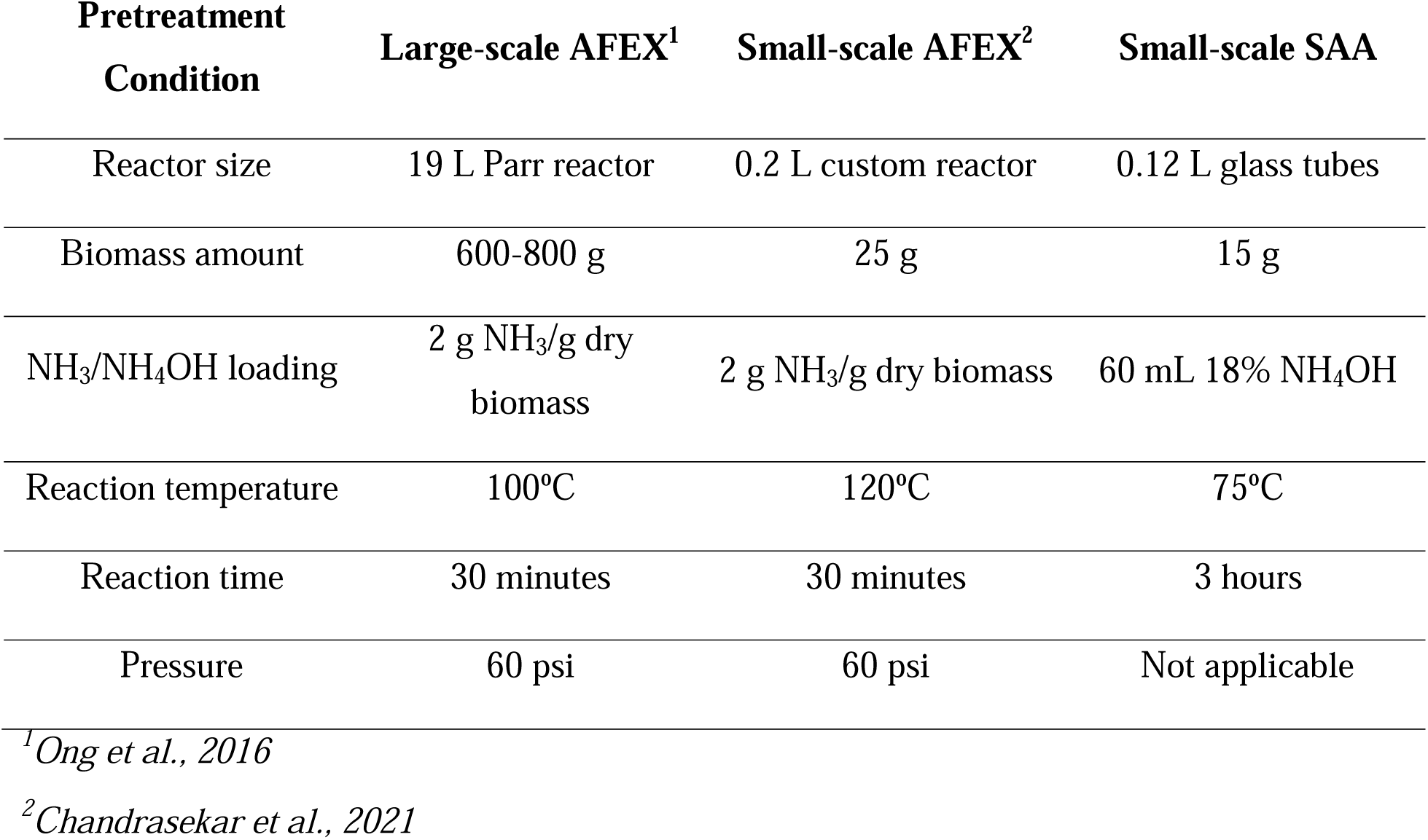
A small-scale Soaking in Aqueous Ammonia pretreatment pipeline generates switchgrass hydrolysates with sugar titers comparable to AFEX pretreatment. (**A**) A schematic diagram of the soaking in aqueous ammonia (SAA) deconstruction pipeline that incorporates enzymatic hydrolysis and microbial fermentation. Comparison of glucose (**B**) and xylose (**C**) titers in hydrolysates generated by AFEX and SAA pretreatment and enzymatic hydrolysis. Columns and bars show the mean and standard deviation (SD) of six independent hydrolysate batches

**Table 1.**
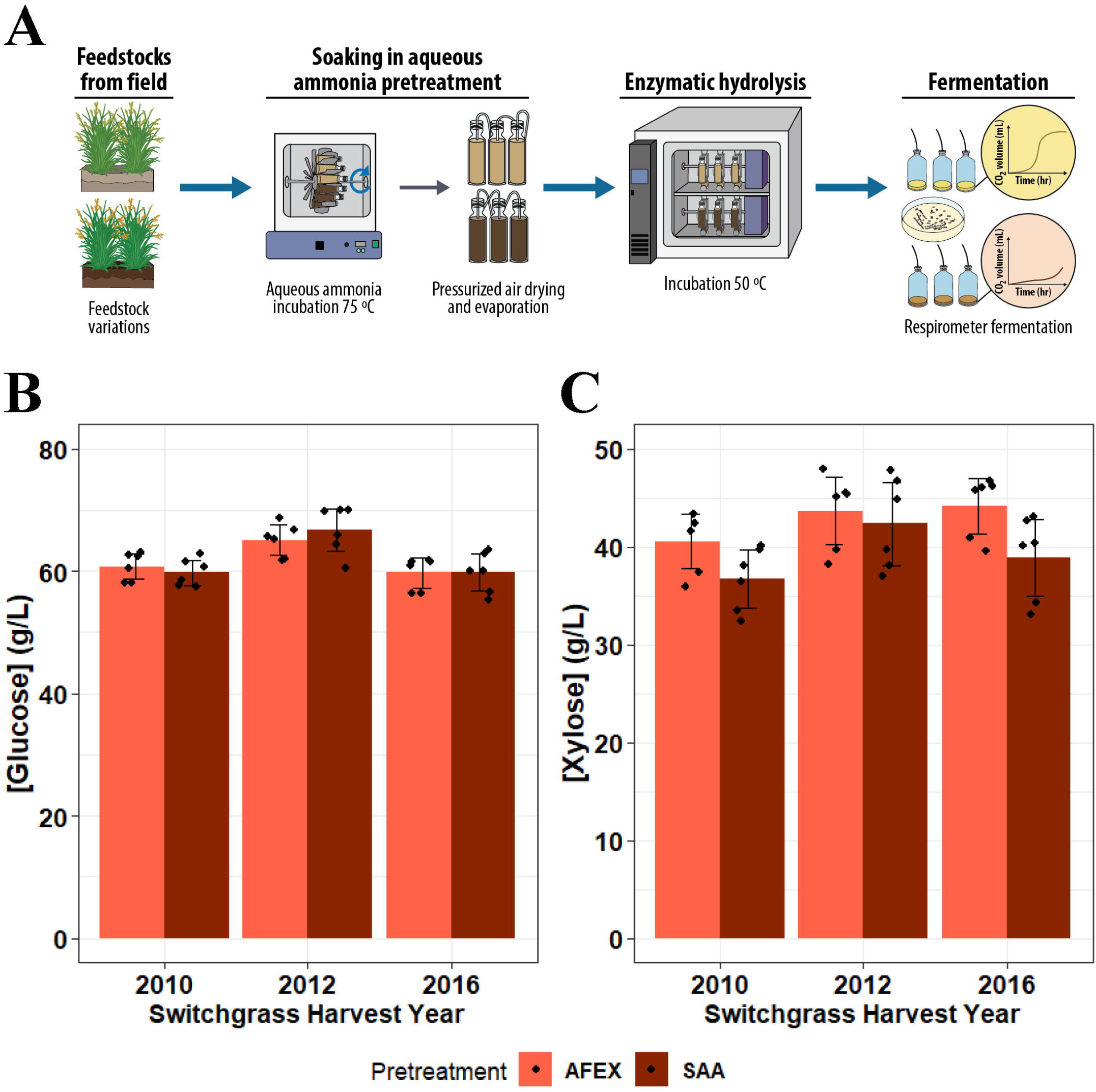
Summary of conditions used for pretreatment 2010, 2012 and 2016 switchgrass.

After hydrolysis of the pretreated biomass, we analyzed the resulting sugar titers to compare the hydrolysis efficiencies between the two different pretreatments. Glucose titers from SAA- and AFEX-deconstructed switchgrass were virtually indistinguishable within harvest years (**Fig. 1B**). Xylose titers were, on average, 3.5 g/L lower (min *p*=0.065, two-sided Wilcoxon tests) across harvest years in SAA-pretreated hydrolysates compared to the paired AFEX-pretreated switchgrass samples (**Fig. 1C**). These results indicated that the SAA pretreatment method generates hydrolysates with similar sugar titers to the AFEX pretreatment method.

To investigate whether different pretreatment methods yielded hydrolysates with distinct inhibitor compositions, we performed targeted quantification of 48 analytes by mass spectrometry (MS) from both SAA- and AFEX-pretreated switchgrass hydrolysates adjusted to pH 5.0 and pH 5.8, except for 2016 at pH 5.0 (**Fig. S1, File S1**). Hydrolysates from each harvest year and pretreatment clustered tightly together in a principal component analysis (PCA) of their chemical compositions, with clear separation between these groups (**Fig. 2A**). The most variance was described by pretreatment method (principal component 1), whereas principal component 2 captured separation by harvest year and precipitation. We next focused on the effect of pretreatment on the concentrations of several characterized microbial inhibitors (**Fig. 2B-C**). As observed previously, AFEX-pretreated switchgrass hydrolysates contained high levels of phenolic amides, such as feruloyl amide and courmaroyl amide. In contrast, the levels of feruloyl amide and coumaroyl amide were approximately 1.4- to 1.8-fold lower in the SAA-deconstructed samples, while the levels of ferulic acid and *p*-coumaric acid were approximately 2.9- to 3.8-fold higher (*p*<0.001, two-sided Wilcoxon tests). There was a strong inverse correlation between the concentrations of coumaric acid and coumaroyl amide (**Fig. 2B**; Spearman’s rho: -0.7, *p*=1.7e-7), as well as ferulic acid and feruloyl amide (**Fig. 3B**; Spearman’s rho: -0.51, *p*=5.3e-4). This result supports the hypothesis that the amides are likely produced by ammonolysis of carboxylic acid groups during AFEX pretreatment (Chundawat et al., 2010). Overall, SAA pretreatment yields hydrolysates with similar sugar content, but they have distinct chemical profiles compared to AFEX pretreatment.

**Fig. 2.**
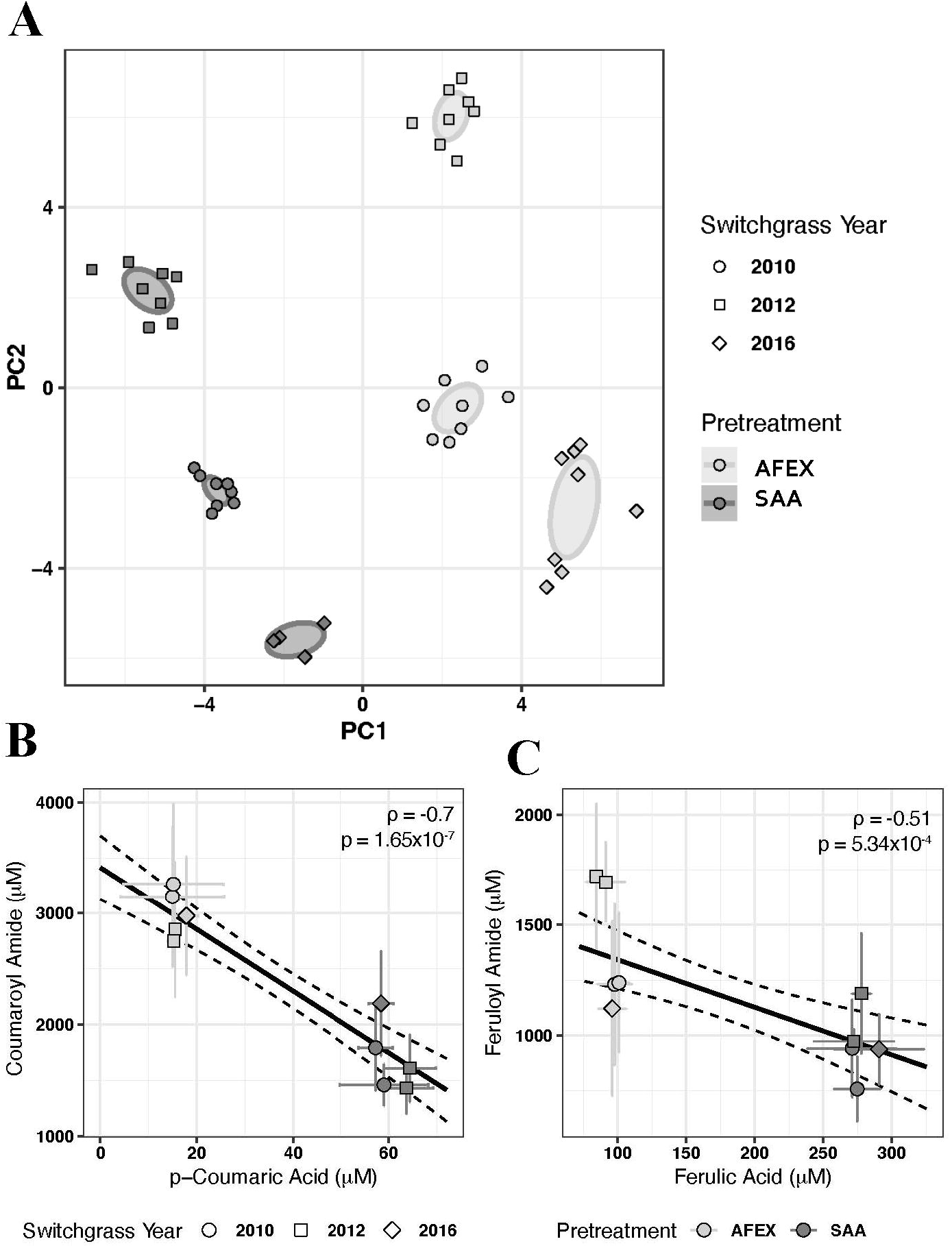
Switchgrass hydrolysates from SAA and AFEX pretreatments are chemically distinct. Principal component analysis (PCA) plot (**A**) displays all the analyzed hydrolysate samples in principal component space for principal components 1 and 2. Ellipses show confidence intervals for each specified group. Scatterplots display the mean and standard deviation of coumaroyl amide and *p*-coumaric acid (**B**), as well as feruloyl amide and ferulic acid (**C**) concentrations from four independent batches of the indicated hydrolysates. Black lines show linear regressions and 95% confidence intervals.

**Figure 3.**
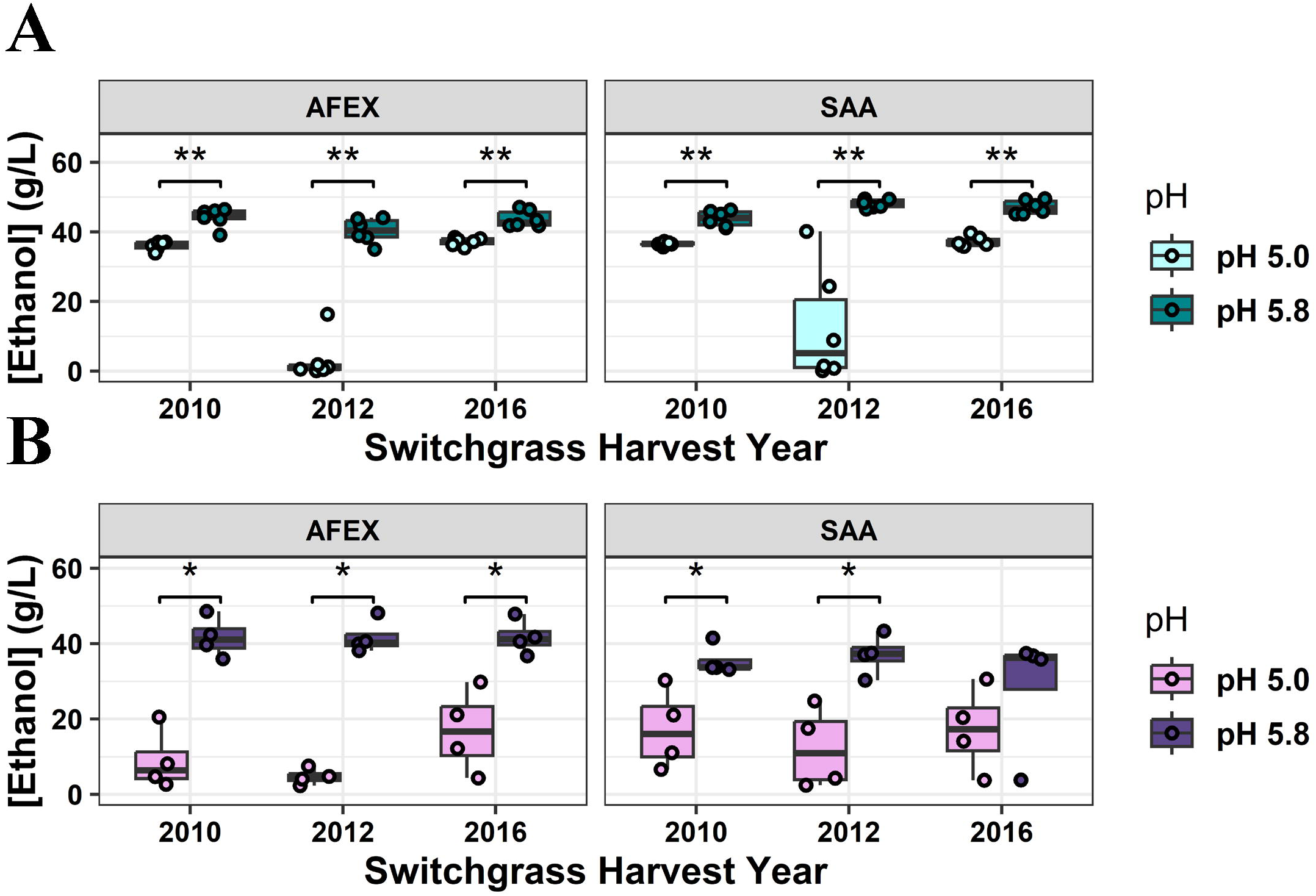
Increasing pH improves yeast and bacterial fermentation in both SAA- and AFEX-pretreated switchgrass hydrolysates. Boxplots display the final ethanol titers from fermentations of switchgrass hydrolysates by the *S. cerevisiae* strain Y1455 (**A**) and the *Z. mobilis* strain Zm2032 (**B**). Brackets denote significance as determined by two-sided Wilcoxon tests; * p<0.05, **p<0.01, ***p<0.001.

### 3.2 Soaking in aqueous ammonia pretreatment modestly relieves drought hydrolysate inhibition

We next tested whether the differing chemical compositions of SAA- and AFEX-pretreated hydrolysates translated into different effects on microbial fermentation at pH of 5.0, particularly for the inhibitory 2012 feedstock. After enzymatic hydrolysis of the pretreated biomass the resulting hydrolysate has pH of approximately 5.0. SAA- and AFEX-pretreated hydrolysates at pH 5.0 were fermented by the engineered, xylose-fermenting *Saccharomyces cerevisiae* strain GLBRCY1455 (see Materials and Methods), as well as by *Zymomonas mobilis* strain Zm2032 (Yang et al., 2018) (**Fig. 3A-B**). We measured final CO_2_ production, lag time (length of delay in minutes for the start of CO_2_ production), glucose and xylose consumption, final cell density (OD_600_), and ethanol titers after 48 hours.

We found fermentations at pH 5.0 of SAA-pretreated drought hydrolysates by Y1455 to be highly variable between hydrolysate batches, with ethanol titers after 48 hours ranging from 0.11 to 49.55 g/L (**Fig. S2A**). These results nonetheless represented an approximately 6-fold increase in median ethanol titers compared to paired AFEX-pretreated hydrolysates, although this difference was not statistically significant (*p*=0.31, two-sided Wilcoxon test). By contrast, no meaningful difference in ethanol production was observed for Y1455 in fermentations of AFEX- and SAA-pretreated hydrolysates from normal precipitation years (**Fig. S2A**). Similar trends were observed for Zm2032, with SAA-pretreated drought hydrolysates yielding 2.5-fold higher median ethanol titers than AFEX-pretreated hydrolysates (**Fig**. **S2B**; *p*=0.49, two-sided Wilcoxon test). Thus, the differing makeup of hydrolysates generated from SAA and AFEX pretreatment did not translate into a full alleviation of the inhibitory effects of drought-grown feedstocks at the pH of 5.0. Moreover, SAA-pretreated drought hydrolysates yielded unpredictable fermentation performances that were hindered by strong batch effects, although still improving ethanol production by Y1455 and Zm2032 on average.

### 3.3 Increasing hydrolysate pH from 5.0 to 5.8 results in increased ethanol production by *S. cerevisiae* and *Z. mobilis*

We next tested the effect of altering hydrolysate pH on fermentation by Y1455 and Zm2023 (**Fig. 3**). Previous research has shown, depending on media conditions, such as temperature, a pH of 4.0-6.0 is optimal for *S. cerevisiae* (Narendranath & Power, 2005), whereas a pH of 5.0-7.5 is optimal for *Z. mobilis* (Lawford et al., 1988). Due to these inherent differences in pH preferences, previous comparisons between these microbes have often been made using different pH values that were tailored to the microbe’s preference under optimal non-hydrolysate conditions (Ong et al., 2016; Zhang et al., 2018). Across both pretreatment methods and all feedstock years, pH adjustment to 5.8 increased median ethanol production by Y1455 by a minimum of 12.2% (*p*=0.002, two-sided Wilcoxon test). The greatest increase in 48-hour ethanol production by Y1455 was observed in AFEX-pretreated drought switchgrass hydrolysate, where median ethanol titers leapt approximately 46-fold, from 0.88 g/L at pH 5.0 to 40.39 g/L at pH 5.8 (**Fig. 3A**). A similar trend observed for fermentations by Zm2032, where median ethanol production at pH 5.8 was an average of 4.26-fold higher than in matched hydrolysates at pH 5.0 (**Fig. 3B**). Here, as well, the greatest increase of ∼9-fold was observed for AFEX-pretreated hydrolysates from drought switchgrass.

Similar trends were observed for CO_2_ production (**Fig. S3**), lag phase (**Fig. S4**), cell growth (**Fig. S5**), and glucose and xylose consumption (**Fig. S6-7**). Y1455 produced significantly more CO_2_ at pH 5.8 than at pH 5.0 within pretreatment types for all three harvest year hydrolysates (all *p*≤0.005, two-sided Wilcoxon tests; **Fig. S3A**). Similarly, Zm2032 produced significantly lower final CO_2_ at pH 5.0 than at pH 5.8 for the hydrolysates from each pretreatment method (**Fig. S3B**), with the exception of the 2016 SAA-pretreated hydrolysates (*p*=0.34, two-sided Wilcoxon test). Lag time was generally shorter for pH 5.8 fermentations by Y1455, but it could not be accurately estimated for fermentations that experienced severe inhibition, including all pH 5.0 fermentations by Zm2032 (**Fig. S4**). Final cell densities (OD_600_) for both Y1455 and Zm2032 were universally higher in pH 5.8 fermentations (**Fig. S5**), although these differences were not statistically significant for some *Z. mobilis* experiments.

Finally, these trends of increased cell growth and fermentation at higher pH were recapitulated in sugar consumption profiles (**Fig. S6-7**). Y1455 nearly fermented all glucose present in hydrolysates from non-drought years, regardless of pH or pretreatment. However, pH adjustment rescued glucose consumption by Y1455 in drought hydrolysates, increasing the median percent consumed by ∼82-fold and ∼5.8-fold in AFEX- and SAA-pretreated hydrolysates, respectively (**Fig. S6A-B**). Y1455 consumed an average of 1.7-fold more xylose from AFEX-pretreated hydrolysates and 2.5-fold more xylose from SAA-pretreated hydrolysates at pH 5.8 compared to pH 5.0 (all *p*≤0.004, two-sided Wilcoxon tests; **Fig. S6C-D**). Zm2032 also nearly completely fermented all glucose in pH 5.8 hydrolysates from all years, regardless of pretreatment, (all *p*≤0.03 except 2016 SAA, two-sided Wilcoxon tests; **Fig. S7A-B**). Zm2032 consumed ∼1.9-fold more xylose from AFEX-pretreated hydrolysates and ∼1.3-fold more from SAA-pretreated hydrolysates at pH 5.8 (all *p*=0.03 except 2016 SAA, two-sided Wilcoxon tests; **Fig. S7C-D).** Therefore, pH adjustment afforded substantial improvements to ethanol production and fermentation by *S. cerevisiae* and *Z. mobilis* across hydrolysates with varying chemical compositions and inhibitory effects.

### 3.4 Ethanol production by *S. cerevisiae* and *Z. mobilis* is greater at higher pH

Our results suggested that pH adjustment might represent a generalizable intervention to alleviate the effects of lignocellulose-derived (LCD) inhibitors in diverse hydrolysates. To test this hypothesis, we developed synthetic mimics of AFEX-pretreated switchgrass hydrolysate (Synthetic Hydrolysate or “SynH”). Previously, we described versions of SynH that were constructed from pure compounds that were identified and quantified from AFEX-pretreated corn stover hydrolysate (Keating et al., 2014). For this study, we utilized compositional data from chemical analysis of AFEX-pretreated switchgrass hydrolysate (Zhang et al., 2020) to formulate a synthetic version of switchgrass hydrolysate. Furthermore, because SynH is reconstituted with pure, off-the-shelf chemical compounds, we generated SynH without (termed “SynHv4.0”) and with many, but not all, of LCD inhibitors (termed “SynHv4.1”), including ferulic acid and *p*-coumaric acid, as well as and feruloyl amide and coumaroyl amide.

To test the interaction between pH and LCD inhibitory compounds on yeast and bacterial fermentations, each engineered *S. cerevisiae* and *Z. mobilis* strain was cultured in SynHv4.0 or SynHv4.1 media at pH 3, 4, 5, 6, and 7, and final ethanol titers after 48 hours were quantified. We observed clear dose-response kinetics between pH and ethanol production (**Fig. 4**), with notable differences between microorganisms and SynH formulations. Ethanol production was universally inhibited at pH 3, and in SynHv4.1 at pH 4. Both microbes produced substantially more ethanol from the LCD inhibitor-less SynHv4.0 than SynHv4.1 at pH 4, indicating a key interaction between pH and the action of LCD inhibitors. Supporting this notion, there were no statistically significant differences in ethanol production from SynHv4.0 and SynHv4.1 by either strain between pH 5 and 7. The lack of difference at pH 5 compared to higher pH levels is likely due to missing inhibitors in the synthetic hydrolysates that are found in switchgrass hydrolysate. Increasing pH between 4 and 7 generally led to increasing ethanol titers by both strains and in both SynH formulations, with notable differences in the inflection points of non-linear trends. For example, Y1455 produced only ∼2.8% more ethanol from SynHv4.0 at pH 5 compared to pH 4, whereas Zm2032 produced ∼3.1-fold more (**Fig. 4**). These differences likely reflect each organism’s preferred pH range (Lawford et al., 1988; Narendranath & Power, 2005), in the absence of perturbations by LCD inhibitors. Between pH 5 and pH 4 in SynHv4.1, Y1455 ethanol titers increased ∼12-fold, and Zm2032 ethanol titers increased ∼17-fold. Between pH 5 and pH 7 in both SynH types, we further observed substantial linear increases in ethanol production by both microbes, with median titers ∼24% higher for Y1455 (adjusted R^2^: 0.72, *p*=1e^-7^, linear regression) and ∼9% higher for Zm2023 (adjusted R^2^: 0.09, *p*=0.12, linear regression). Finally, the effect of pH on ethanol production was recapitulated in overall fermentative growth as measured by CO_2_ production (**Fig. S8**), cell density (**Fig. S9**), and sugar consumption (**Fig. S10**). Thus, increasing pH to the range of 5–7 greatly improved both growth and metabolism by Y1455 and Zm2032, especially in the presence of LCD inhibitors.

**Figure 4.**
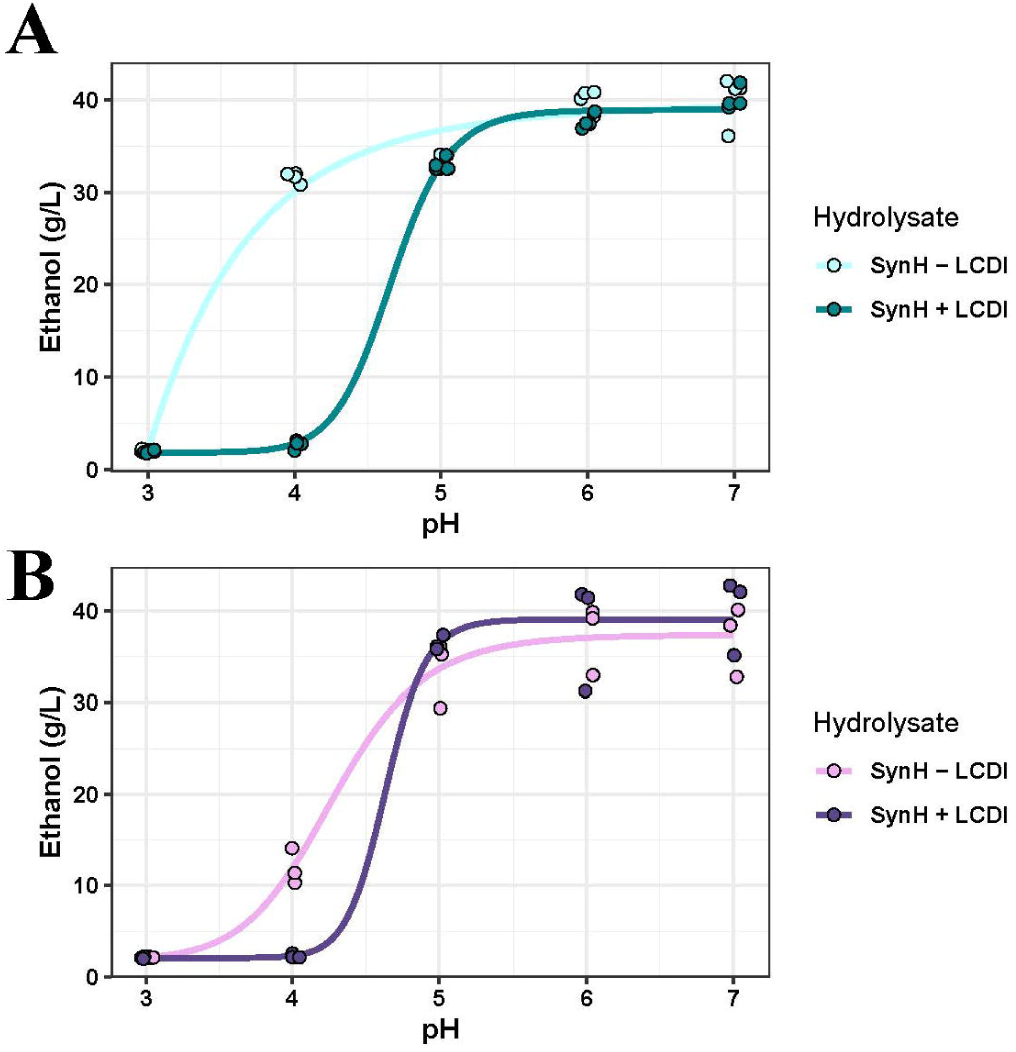
Increasing pH of synthetic hydrolysates increases ethanol production by *S. cerevisiae* and *Z. mobilis*. Dose-response curves display final ethanol titers from fermentations of synthetic hydrolysate without (SynH - LCDI, SynHv4.0) or with lignocellulose-derived inhibitors (SynH + LCDI, SynHv4.1) at pH 3-7 by *S. cerevisiae* Y1455 (**A**) and *Z. mobilis* Zm2032 (**B**).

### 3.5 Increasing hydrolysate pH increases isobutanol titers by hybrid *S. cerevisiae* ethanol and isobutanol co-producing strain

Isobutanol (IBA) is a potential substrate for the formation of alkanes and alkenes used for aviation fuel production, making it a key second-generation biofuel (Geleynse et al., 2018; Holladay et al., 2020). The engineered hybrid *S. cerevisiae* strain yHRW253 (Pastore de Lima et al., 2023), which can simultaneously ferment sugars from hydrolysate into ethanol and isobutanol, also experiences inhibition by drought hydrolysates. We therefore tested whether isobutanol production by this strain could also be improved by increasing the hydrolysate pH. We performed fermentation experiments with yHRW253 in both AFEX-pretreated switchgrass hydrolysate and SAA-pretreated switchgrass hydrolysate at pH 5.0 and 5.8.

Neither pretreatment nor pH had a significant effect on lag phase (**Fig. S12A-B**) or total cell growth (**Fig. S12C-D**) for yHRW253. CO_2_ production was only significantly different, lower in this case, in SAA-pretreated 2016 hydrolysate (*p*=0.03; **Fig. S12E-F**). Glucose was nearly completely consumed in all hydrolysates from normal precipitation years at pH 5.0 and 5.8 (**Fig. S12G-H**). Increasing the pH of the drought year switchgrass hydrolysates recapitulated the near total consumption of glucose seen in normal year switchgrass hydrolysates, but not in a statistically significant manner (*p*≥0.3, two-sided Wilcoxon tests).

Median ethanol production was modestly lower (∼19% on average) at pH 5.8 than pH 5.0 across all feedstock years and deconstruction methods (**Fig. 5A**), although this difference was only statistically significant in the 2010 SAA-pretreated hydrolysate (*p*=0.03, two-sided Wilcoxon test). Interestingly, comparisons between pretreatment methods of the same harvest year switchgrass at pH 5.0 and at 5.8 did not have a significant effect on ethanol production (min *p*=0.06, two-sided Wilcoxon tests, **Fig. S11**). Crucially, almost no isobutanol was produced from 2012 hydrolysates at pH 5.0 (**Fig. 5B**). In 2012 AFEX- and SAA-pretreated hydrolysates, and in 2010 SAA-pretreated hydrolysate, there were statistically significant increases in isobutanol production at pH 5.8 compared to 5.0 (max *p*=0.03, two-sided Wilcoxon tests) (**Fig. 5B**). Notably, fermentation of SAA- and AFEX-pretreated hydrolysates at pH 5.8 yielded ∼5.9-fold and ∼1.8-fold more isobutanol, respectively, than fermentations of the same hydrolysates at pH 5.0 in non-drought years (**Fig. 5B**). By contrast, for the inhibitory 2012 hydrolysates, median isobutanol production increased in SAA- and AFEX-pretreated hydrolysates at pH 5.8 by ∼30- fold and ∼56-fold respectively. Overall, the quantitative effect of increased pH suggests that pH adjustment is a promising strategy to improve isobutanol production from lignocellulosic hydrolysates by engineered *S. cerevisiae*.

**Figure 5.**
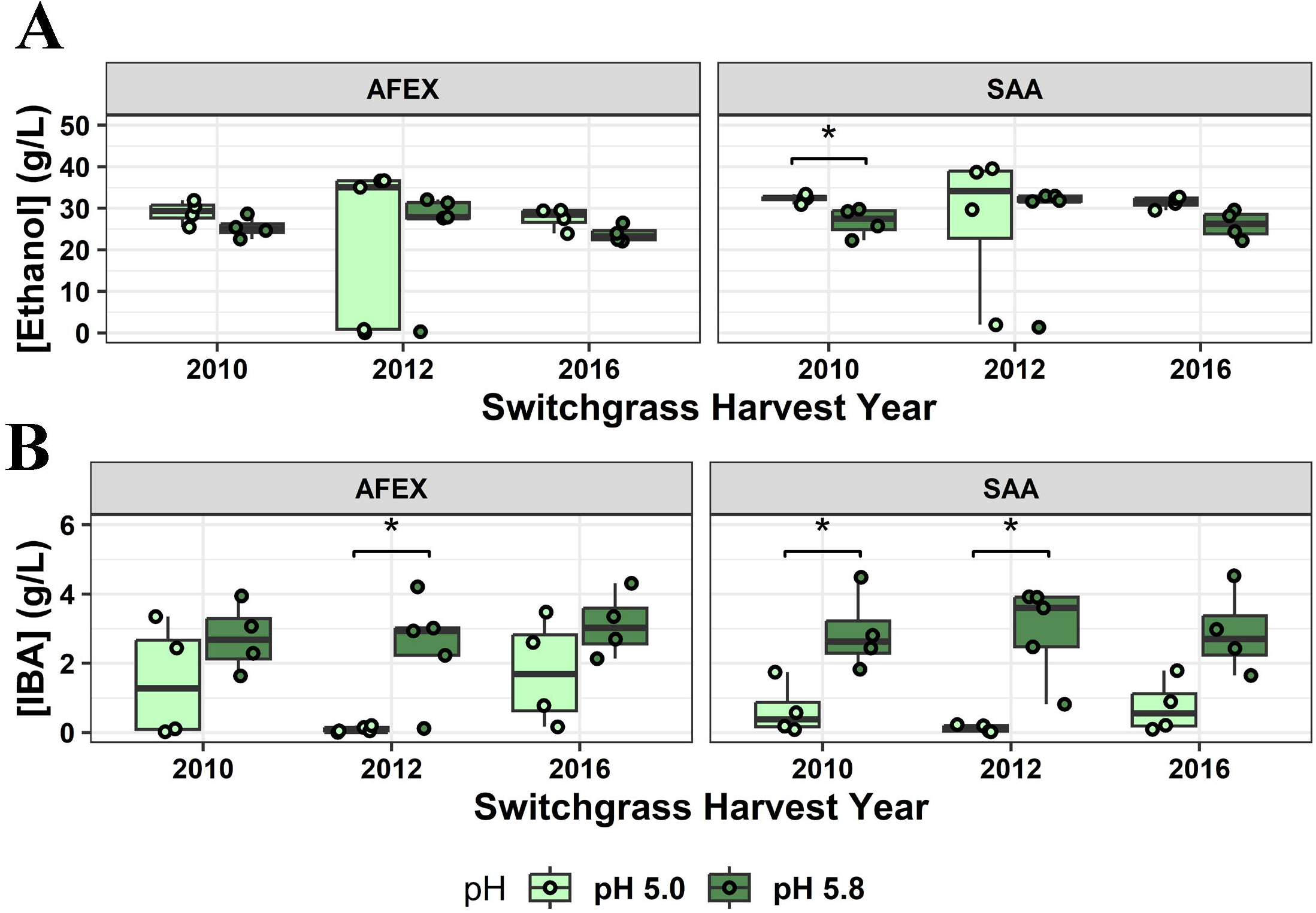
pH adjustment improves isobutanol (IBA) production from hydrolysates. Boxplots display final ethanol (**A**) and isobutanol (**B**) titers from SAA- and AFEX-pretreated hydrolysates from three harvest years, fermented at pH 5.0 and 5.8 by *S. cerevisiae* yHRW253 hybrid. Brackets denote significance level as determined by two-sided Wilcoxon tests; * p<0.05, **p<0.01, ***p<0.001.

## 4. Discussion

To identify ways to overcome the impaired fermentation of drought switchgrass by engineered *S. cerevisiae* and *Z. mobilis*, we compared the fermentations of drought and non-drought switchgrass hydrolysates at two different pH values, and from two different pretreatment methods. We first tested whether the inhibitory effect of hydrolysates from drought year feedstocks could be alleviated by using the SAA pretreatment method, which produced similar yields of glucose and xylose compared to the AFEX-pretreatment (**Fig. 1B-C)**. The overall composition of the SAA-pretreated and AFEX-pretreated hydrolysates differed greatly, including increased concentrations of inhibitory amides in AFEX-pretreated hydrolysates compared to higher concentrations of the conjugate acids in SAA-pretreated hydrolysates (**Fig. 2**). Phenolic acids such as ferulic acid, *p*-coumaric acid, and other compounds can inhibit *S. cerevisiae* fermentation (Adeboye et al., 2014; Klinke et al., 2004; Mussatto & Roberto, 2004), as can high levels of acetic acid (Palmqvist et al., 2000). *Z. mobilis* also experiences inhibitory effects from phenolic compounds found in lignocellulosic-derived hydrolysates (Franden et al., 2013). We found ferulic acid and *p*-coumaric acid were at higher concentrations in SAA-pretreated hydrolysate, whereas feruloyl amide and coumaroyl amide were at higher concentrations in AFEX-pretreated hydrolysate, producing an inverse relationship between pretreatment and acid vs. amide formation (**Fig. 2B-C**). This difference is likely due to the SAA pretreatment leading to lower conversion of the acids to their amide forms than AFEX pretreatment from ammonolysis reactions (Chundawat et al., 2010). A benefit of the SAA pretreatment is that it uses lower temperature, pressure, and biomass, which could create a lower energy cost for the process (**Table 1**). SAA pretreatment should also have lower capital costs, since the AFEX pretreatment needs specialized reactors to safely accommodate the high pressures used with ammonia gas. This could lead to a more resilient and economical process. Therefore, the SAA pretreatment method is comparable to AFEX pretreatment for the deconstruction of switchgrass and potentially more cost effective.

As SAA-pretreatment alone did not relieve the inhibitory effects of drought-grown switchgrass hydrolysates on microbial fermentation, we next tested the effect of manipulating hydrolysate pH. Increased pH has been previously shown to increase ethanol production by yeast (Narendranath & Power, 2005) through mechanisms that include the neutralization of weak acids (Lam et al., 2021), and increased pH has also been shown to increase ethanol production by *Z. mobilis* (Lawford et al., 1988). The higher level of acids may render SAA-pretreated hydrolysates more amenable to intervention, because pH adjustment from 5.0 to 5.8 can shift more of the weak acids into their deprotonated forms (ferulic acid pKa = 4.58; *p*-coumaric acid pKa = 4.64), preventing them for passively diffusing into the cell and acidifying the cytosol, thereby mitigating their inhibitory effects. In contrast the pH adjustment from 5.0 to 5.8 has little effect on the basic amides within this pH range. This led us to believe that the microbes in the SAA-pretreated hydrolysate at pH 5.8 grows to higher cell density and produces more ethanol or isobutanol (**Fig. 3, Fig. S5, Fig. S10-11**) because the neutralization of acids in the media created a less inhibitory environment for fermentation and growth. This result does not explain overall increases in ethanol or isobutanol yields at higher pH AFEX-pretreated hydrolysates, and further study into the tradeoffs between producing these competing alcohol products is needed. The increased pH could still have neutralization effects on the lower concentrations of acids within AFEX-pretreated hydrolysates by creating a less inhibitory environment. A hydrolysate pH of 5.8 could be nearer to the optimal pH value for these microbes to grow in the presence of LCD inhibitors (**Fig. 4**). Overall, we found that the most promising method to overcome the inhibitory effect of hydrolysates from drought-affected switchgrass on downstream microbial fermentation was to increase the pH, regardless of the pretreatment method. We show that this intervention is sufficient to substantially increase production of biofuels by strains of *S. cerevisiae* and *Z. mobilis* from both real and synthetic hydrolysates with diverse chemical compositions.

## 5. Conclusion

We tested adjustments to hydrolysate composition in an effort to develop sustainable and economical methods to create lignocellulosic-derived biofuels from switchgrass, while acknowledging the influence on microbial fermentation of uncontrollable factors during plant growth. Due to climate change, extreme weather events, such as drought, are predicted to become more frequent. Droughts, such as in Wisconsin in 2012, can affect the composition of biofuel crops like switchgrass, which may in turn impact microbial biofuel production by lignocellulosic biorefineries. Previous work has shown the inhibitory effect of drought year switchgrass hydrolysate on *S. cerevisiae* fermentation, and we have shown that increased pH could be a method to overcome inhibition seen from fermenting 2012 switchgrass hydrolysate. This work shows that the selection of pretreatment methods and the pH of deconstructed sugar streams are important factors in producing sustainable biofuels from switchgrass grown under different environmental conditions.

## Acknowledgments

We dedicate this manuscript to the late Dr. Yaoping Zhang, a dedicated and innovative scientist who will be greatly missed. This material is based upon work supported by the Great Lakes Bioenergy Research Center, U.S. Department of Energy, Office of Science, Biological and Environmental Research Program under Award Number DE-SC0018409. Research in the Hittinger Lab is further supported by the National Science Foundation under Grant No. DEB-2110403, the USDA National Institute of Food and Agriculture (Hatch Project 7005101), and an H. I. Romnes Faculty Fellowship, supported by the Office of the Vice Chancellor for Research and Graduate Education with funding from the Wisconsin Alumni Research Foundation, respectively. J.G.C. was supported by a Predoctoral Training Grant in Genetics funded by the National Institutes of Health under Grant No. T32GM007133 and by the National Science Foundation Graduate Research Fellowship Program under Grant No. DGE-1747503. We thank Kurt Creamer and Novozymes for generously providing enzymes, the GLBRC Metabolomics Facility for quantification of analytes, Matt Wisniewski and Chelsea Mamott for graphics production, Dr. Larry Anthony, Butamax and DuPont R&D for their contributions to the yHRW253 hybrid yeast strain, Dr. Troy Runge, and Dr. Rebecca Garlock Ong and members of the Sato and Hittinger labs for helpful suggestions. Graphical abstract was created in BioRender, Sato, T. (2025) https://BioRender.com/l22d704 (Agreement number: QQ27RX43JW).

## References

Adeboye, P. T., Bettiga, M., & Olsson, L. (2014). The chemical nature of phenolic compounds determines their toxicity and induces distinct physiological responses in Saccharomyces cerevisiae in lignocellulose hydrolysates. AMB Express, 4(1), 1–10. 10.1186/s13568-014-0046-7

Alcázar, M., Kind, T., Gschaedler, A., Silveria, M., Arrizon, J., Fiehn, O., Vallejo, A., Higuera, I., & Lugo, E. (2017). Effect of steroidal saponins from Agave on the polysaccharide cell wall composition of Saccharomyces cerevisiae and Kluyveromyces marxianus. LWT, 77, 430–439. 10.1016/J.LWT.2016.11.048

Chandrasekar, M., Joshi, L., Krieg, K., Chipkar, S., Burke, E., Debrauske, D. J., Thelen, K. D., Sato, T. K., & Ong, R. G. (2021). A high solids field-to-fuel research pipeline to identify interactions between feedstocks and biofuel production. Biotechnology for Biofuels, 14(1). 10.1186/s13068-021-02033-6

Chipkar, S., Smith, K., Whelan, E. M., Debrauske, D. J., Jen, A., Overmyer, K. A., Senyk, A., Hooker-Moericke, L., Gallmeyer, M., Coon, J. J., Jones, A. D., Sato, T. K., & Ong, R. G. (2022). Water-soluble saponins accumulate in drought-stressed switchgrass and may inhibit yeast growth during bioethanol production. Biotechnology for Biofuels and Bioproducts, 15(1), 116. 10.1186/s13068-022-02213-y

Chundawat, S. P. S., Vismeh, R., Sharma, L. N., Humpula, J. F., da Costa Sousa, L., Chambliss, C. K., Jones, A. D., Balan, V., & Dale, B. E. (2010). Multifaceted characterization of cell wall decomposition products formed during ammonia fiber expansion (AFEX) and dilute acid based pretreatments. Bioresource Technology, 101(21), 8429–8438. 10.1016/j.biortech.2010.06.027

Dunson Jr., J. B., Tucker III, M. P., Elander, R. T., & Hennessey, S. M. (2007). Treatment of biomass to obtain a target chemical (US7781191B2). U.S. Patent and Trademark Office. https://patentimages.storage.googleapis.com/54/eb/27/18947bf1b17a72/US7781191.pdf

Enright, A. L., Banta, A. B., Ward, R. D., Rivera Vazquez, J., Felczak, M. M., Wolfe, M. B., TerAvest, M. A., Amador-Noguez, D., & Peters, J. M. (2023). The genetics of aerotolerant growth in an alphaproteobacterium with a naturally reduced genome. MBio, 14(6). 10.1128/mbio.01487-23

Franden, M. A., Pilath, H. M., Mohagheghi, A., Pienkos, P. T., & Zhang, M. (2013). Inhibition of growth of Zymomonas mobilis by model compounds found in lignocellulosic hydrolysates. Biotechnology for Biofuels, 6(1). 10.1186/1754-6834-6-99

Gambacorta, F. V, Wagner, E. R., Jacobson, T. B., Tremaine, M., Muehlbauer, L. K., McGee, M. A., Baerwald, J. J., Wrobel, R. L., Wolters, J. F., Place, M., Dietrich, J. J., Xie, D., Serate, J., Gajbhiye, S., Liu, L., Vang-Smith, M., Coon, J. J., Zhang, Y., Gasch, A. P., … Pfleger, B. F. (2022). Comparative functional genomics identifies an iron-limited bottleneck in a Saccharomyces cerevisiae strain with a cytosolic-localized isobutanol pathway. Synthetic and Systems Biotechnology, 7(2), 738–749. 10.1016/j.synbio.2022.02.007

Geleynse, S., Brandt, K., Garcia-Perez, M., Wolcott, M., & Zhang, X. (2018). The Alcohol-to-Jet Conversion Pathway for Drop-In Biofuels: Techno-Economic Evaluation. ChemSusChem, 11(21), 3728–3741. 10.1002/cssc.201801690

Georgopapadakou, N. H., & Walsh, T. J. (1996). MINIREVIEW Antifungal Agents: Chemotherapeutic Targets and Immunologic Strategies. ANTIMICROBIAL AGENTS AND CHEMOTHERAPY, 40(2), 279–291. https://journals.asm.org/journal/aac

Güldener, U., Heck, S., Fielder, T., Beinhauer, J., & Hegemann, J. H. (1996). A new efficient gene disruption cassette for repeated use in budding yeast. Nucleic Acids Research, 24(13), 2519–2524. 10.1093/nar/24.13.2519

Holladay, J., Abdullah, Z., & Heyne, J. (2020). Sustainable Aviation Fuel: Review of Technical Pathways Report.

Jit, I., And, S., & Feingold, D. S. (1981). Heterogeneity of Action Mechanisms Among Antimycotic Imidazoles. In ANTIMICROBIAL AGENTS AND CHEMOTHERAPY (Vol. 20, Issue 1). https://journals.asm.org/journal/aac

Keating, D. H., Zhang, Y., Ong, I. M., McIlwain, S., Morales, E. H., Grass, J. A., Tremaine, M., Bothfeld, W., Higbee, A., Ulbrich, A., Balloon, A. J., Westphall, M. S., Aldrich, J., Lipton, M. S., Kim, J., Moskvin, O. V., Bukhman, Y. V., Coon, J. J., Kiley, P. J., … Landick, R. (2014). Aromatic inhibitors derived from ammonia-pretreated lignocellulose hinder bacterial ethanologenesis by activating regulatory circuits controlling inhibitor efflux and detoxification. Frontiers in Microbiology, 5. 10.3389/fmicb.2014.00402

Keshwani, D. R., & Cheng, J. J. (2009). Switchgrass for bioethanol and other value-added applications: A review. In Bioresource Technology (Vol. 100, Issue 4, pp. 1515–1523). 10.1016/j.biortech.2008.09.035

Kim, T. H., & Lee, Y. Y. (2007). Pretreatment of corn stover by soaking in aqueous ammonia at moderate temperatures. Applied Biochemistry and Biotechnology, 137–140(1–12), 81–92. 10.1007/s12010-007-9041-7

Klinke, H. B., Thomsen, A. B., & Ahring, B. K. (2004). Inhibition of ethanol-producing yeast and bacteria by degradation products produced during pre-treatment of biomass. In Applied Microbiology and Biotechnology (Vol. 66, Issue 1, pp. 10–26). 10.1007/s00253-004-1642-2

Ko, J. K., Bak, J. S., Jung, M. W., Lee, H. J., Choi, I.-G., Kim, T. H., & Kim, K. H. (2009). Ethanol production from rice straw using optimized aqueous-ammonia soaking pretreatment and simultaneous saccharification and fermentation processes. Bioresource Technology, 100(19), 4374– 4380. 10.1016/j.biortech.2009.04.026

Lam, F. H., Turanli-Yildiz, B., Liu, D., Resch, M. G., Fink, G. R., & Stephanopoulos, G. (2021). Engineered yeast tolerance enables efficient production from toxified lignocellulosic feedstocks. Science Advances, 7(26). 10.1126/sciadv.abf7613

Lawford, H., Holloway, P., & Ruggiero, A. (1988). EFFECT OF pH ON GROWTH AND ETHANOL PRODUCTION BY Zymomonas. In Biotechnology Letters (Vol. 0).

Lee, S.-B., Tremaine, M., Place, M., Liu, L., Pier, A., Krause, D. J., Xie, D., Zhang, Y., Landick, R., Gasch, A. P., Hittinger, C. T., & Sato, T. K. (2021). Crabtree/Warburg-like aerobic xylose fermentation by engineered Saccharomyces cerevisiae. Metabolic Engineering, 68, 119–130. 10.1016/j.ymben.2021.09.008

Mallya, G., Zhao, L., Song, X. C., Niyogi, D., & Govindaraju, R. S. (2013). 2012 Midwest Drought in the United States. Journal of Hydrologic Engineering, 18(7), 737–745. 10.1061/(asce)he.1943-5584.0000786

Mugford, S. T., & Osbourn, A. (2013). Saponin synthesis and function. In Isoprenoid Synthesis in Plants and Microorganisms: New Concepts and Experimental Approaches. 10.1007/978-1-4614-4063-5_28

Mussatto, S. I., & Roberto, I. C. (2004). Alternatives for detoxification of diluted-acid lignocellulosic hydrolyzates for use in fermentative processes: A review. In Bioresource Technology (Vol. 93, Issue 1, pp. 1–10). Elsevier Ltd. 10.1016/j.biortech.2003.10.005

Narendranath, N. V., & Power, R. (2005). Relationship between pH and Medium Dissolved Solids in Terms of Growth and Metabolism of Lactobacilli and Saccharomyces cerevisiae during Ethanol Production. Applied and Environmental Microbiology, 71(5), 2239. 10.1128/AEM.71.5.2239-2243.2005

Ong, R. G., Higbee, A., Bottoms, S., Dickinson, Q., Xie, D., Smith, S. A., Serate, J., Pohlmann, E., Jones, A. D., Coon, J. J., Sato, T. K., Sanford, G. R., Eilert, D., Oates, L. G., Piotrowski, J. S., Bates, D. M., Cavalier, D., & Zhang, Y. (2016). Inhibition of microbial biofuel production in drought-stressed switchgrass hydrolysate. Biotechnology for Biofuels, 9, 237. 10.1186/s13068-016-0657-0

Palmqvist, E., Arbel, B., & Agerdal, H.-H. (2000). Fermentation of lignocellulosic hydrolysates. II: inhibitors and mechanisms of inhibition.

Pastore de Lima, A. E., Wrobel, R. L., Paul, B., Anthony, L. C., Sato, T. K., Zhang, Y., Hittinger, C. T., & Maravelias, C. T. (2023). High yield co-production of isobutanol and ethanol from switchgrass: experiments, and process synthesis and analysis. Sustainable Energy and Fuels, 7(14). 10.1039/d2se01741e

Piotrowski, J. S., Zhang, Y., Bates, D. M., Keating, D. H., Sato, T. K., Ong, I. M., & Landick, R. (2014). Death by a thousand cuts: The challenges and diverse landscape of lignocellulosic hydrolysate inhibitors. Frontiers in Microbiology, 5(MAR). 10.3389/fmicb.2014.00090

Pryor, S. W., Karki, B., & Nahar, N. (2011). Enzymatic Hydrolysis of Switchgrass and Tall Wheatgrass Mixtures Using Dilute Sulfuric Acid and Aqueous Ammonia Pretreatments. Biological Engineering, 3(3), 163–171.

Rinehart, L. (2006). A Publication of ATTRA-National Sustainable Agriculture Information Service • 1-800-346-9140 • www.attra.ncat.org. www.attra.ncat.org

Schmer, M. R., Vogel, K. P., Mitchell, R. B., Perrin, R. K., & Matson, P. A. (2008). Net energy of cellulosic ethanol from switchgrass. www.pnas.orgcgidoi10.1073pnas.0704767105

Schwalbach, M. S., Keating, D. H., Tremaine, M., Marner, W. D., Zhang, Y., Bothfeld, W., Higbee, A., Grass, J. A., Cotten, C., Reed, J. L., da Costa Sousa, L., Jin, M., Balan, V., Ellinger, J., Dale, B., Kiley, P. J., & Landick, R. (2012). Complex physiology and compound stress responses during fermentation of alkali-pretreated corn stover hydrolysate by an Escherichia coli ethanologen. Applied and Environmental Microbiology, 78(9), 3442–3457. 10.1128/AEM.07329-11

Serate, J., Xie, D., Pohlmann, E., Donald, C., Shabani, M., Hinchman, L., Higbee, A., Mcgee, M., La Reau, A., Klinger, G. E., Li, S., Myers, C. L., Boone, C., Bates, D. M., Cavalier, D., Eilert, D., Oates, L. G., Sanford, G., Sato, T. K., … Zhang, Y. (2015). Controlling microbial contamination during hydrolysis of AFEX-pretreated corn stover and switchgrass: effects on hydrolysate composition, microbial response and fermentation. Biotechnology for Biofuels, 8(1), 180. 10.1186/s13068-015-0356-2

Sokolov, S. S., Popova, M. M., Pohl, P., Horner, A., Akimov, S. A., Kireeva, N. A., Knorre, D. A., Batishchev, O. V., & Severin, F. F. (2022). Structural Role of Plasma Membrane Sterols in Osmotic Stress Tolerance of Yeast Saccharomyces cerevisiae. Membranes, 12(12). 10.3390/membranes12121278

van der Pol, E. C., Bakker, R. R., Baets, P., & Eggink, G. (2014). By-products resulting from lignocellulose pretreatment and their inhibitory effect on fermentations for (bio)chemicals and fuels. In Applied Microbiology and Biotechnology (Vol. 98, Issue 23). 10.1007/s00253-014-6158-9

Xie, S., Li, Z., Luo, S., & Zhang, W. (2024). Bioethanol to jet fuel: Current status, challenges, and perspectives. Renewable and Sustainable Energy Reviews, 192, 114240. 10.1016/j.rser.2023.114240

Yang, S., Vera, J. M., Grass, J., Savvakis, G., Moskvin, O. V., Yang, Y., McIlwain, S. J., Lyu, Y., Zinonos, I., Hebert, A. S., Coon, J. J., Bates, D. M., Sato, T. K., Brown, S. D., Himmel, M. E., Zhang, M., Landick, R., Pappas, K. M., & Zhang, Y. (2018). Complete genome sequence and the expression pattern of plasmids of the model ethanologen Zymomonas mobilis ZM4 and its xylose-utilizing derivatives 8b and 2032. Biotechnology for Biofuels, 11(1), 125. 10.1186/s13068-018-1116-x

Zhang, Y., Oates, L. G., Serate, J., Xie, D., Pohlmann, E., Bukhman, Y. V., Karlen, S. D., Young, M. K., Higbee, A., Eilert, D., Sanford, G. R., Piotrowski, J. S., Cavalier, D., Ralph, J., Coon, J. J., Sato, T. K., & Ong, R. G. (2018). Diverse lignocellulosic feedstocks can achieve high field-scale ethanol yields while providing flexibility for the biorefinery and landscape-level environmental benefits. GCB Bioenergy, 10(11), 825–840. 10.1111/gcbb.12533

Zhang, Y., Serate, J., Xie, D., Gajbhiye, S., Kulzer, P., Sanford, G., Russell, J. D., McGee, M., Foster, C., Coon, J. J., Landick, R., & Sato, T. K. (2020). Production of hydrolysates from unmilled AFEX-pretreated switchgrass and comparative fermentation with Zymomonas mobilis. Bioresource Technology Reports, 11, 100517. 10.1016/j.biteb.2020.100517

Zhao, C., Shao, Q., & Chundawat, S. P. S. (2020). Recent advances on ammonia-based pretreatments of lignocellulosic biomass. In Bioresource Technology (Vol. 298). Elsevier Ltd. 10.1016/j.biortech.2019.122446

