## Supplemental Tables and Figures for "pH adjustment increases biofuel production from inhibitory switchgrass hydrolysates"

**Supplemental Figures**

### Supplemental Table S1. Chemical composition of SynHv4.0 medium

| **Component** | **Source (Catalog # or CAS #, Vendor)** | **Concentration in mM** |
| --- | --- | --- |
| **Major Sugars, Acids and Organics (mM)** | | |
| D-Glucose |  | 333 |
| D-Xylose |  | 253 |
| Sodium formate | S648-500, Fisher Scientific | 8.5 |
| Sodium nitrate | S343-500, Fisher Scientific | 1.1 |
| Sodium succinate.6H_2_O | S413-500, Fisher Scientific | 0.3 |
| Glycerol | G33-500, Fisher Scientific | 2.9 |
| Sodium acetate | S8750-250G, Fisher Scientific | 44.2 |
| D-Mannose | 3458-28-4, Acros | 2.2 |
| L-Arabinose | 28697-53-2, Acros | 5.9 |
| D-Fructose | L96-500, Fisher Scientific | 11.9 |
| D-Galactose | G0750-500G, Sigma Aldrich | 2.2 |
| D-Ribose | 132361000, Acros | 1.3 |
| Acetamide | 00160, Fluka | 53.1 |
| **Abundant Metals and Inorganics (mM)** | | |
| (NH_4_)_2_SO_4_ | 7783-20-2, Acros | 3 |
| NH_4_Cl | A661-500, Fisher Scientific | 24 |
| MgCl_2_·6H_2_O | M33-500, Fisher Scientific | 13.3 |
| CaCl_2_·2H_2_O | C79-500, Fisher Scientific | 1.2 |
| KCl | P217-500, Fisher Scientific | 30.4 |
| NaCl  KH_2_PO_4_ (as a buffer)  K_2_HPO_4_ (as a buffer) | S271-500, Fisher Scientific  P386-500, Fisher Scientific  P288-500, Fisher Scientific | 0.8  4.1  7.7 |
| **Trace Metals, Inorganics, Vitamins and other compounds (μM)** | | |
| ZnCl_2_ | 96469, Fluka | 10 |
| MnCl_2_·4H_2_O | M87-500, Fisher Scientific | 64 |
| CuCl_2_·2H_2_O | 307483-100G, Sigma Aldrich | 1.5 |
| CoCl_2_·6H_2_O | 60820, Fluka | 0.02 |
| H_3_BO_3_ | B6768-500G, Sigma Aldrich | 29.3 |
| (NH_4_)_6_Mo_7_O_24_•4H_2_O | 09878, Fluka | 0.26 |
| Fe (III) citrate·4H_2_O | F6129-250g, Sigma Aldrich | 6.7 |
| Sodium citrate | BP327-500, Fisher Scientific | 1000 |
| Pyridoxine.HCl | BP267750, Fisher Scientific | 2.14 |
| Nicotinic Acid | 1282950000, Acros | 26.78 |
| Biotin | B4501-10G, Sigma Aldrich | 0.1 |
| Myo-Inositol  Thiamine HCl  Calcium Pantothenate  Betaine.H_2_O  Choline Chloride  D/L-Carnitine.HCl | 57570-25G, Fluka  BP892-100, Sigma Aldrich  C8731-25G, Sigma Aldrich  14300, Fluka  67-48-1, Acros  461-05-2, Acros | 56  0.4  3  700  300  300 |
| **Amino Acids and Nucleotides (μM)** | | |
| L-Alanine | 05129-25G, Fluka | 381 |
| L-Arginine.HCl | 11039, Sigma Aldrich | 108 |
| L-Asparagine | 11149-25G-F, Sigma Aldrich | 113 |
| D/L-Aspartic acid.K | A2025-100G, Sigma Aldrich | 195 |
| L-Cysteine.HCl | 30129-25G, Sigma Aldrich | 50 |
| L-Glutamine | 49419-25G, Sigma Aldrich | 76 |
| L-Glutamic acid.K  Glycine | G1501-500G, Sigma Aldrich  410225-250G, Sigma Aldrich | 155  296 |
| L-Histidine | 71-00-1, Acros | 16 |
| L-Isoleucine | 58879, Sigma Aldrich | 59 |
| L-Leucine | 61819-25G, Sigma Aldrich | 229 |
| L-Lysine.HCl | L5626-100G, Sigma Aldrich | 79 |
| L-Methionine | 64319-25G-F, Sigma Aldrich | 100 |
| L-Phenylalanine | P5482-25G, Sigma Aldrich | 207 |
| L-Proline | 81709-25G, Sigma Aldrich | 141 |
| L-Serine | 84959-100G, Sigma Aldrich | 144 |
| L-Threonine | 89179-10G, Sigma Aldrich | 100 |
| L-Tryptophan | 93659-10G, Sigma Aldrich | 50 |
| L-Tyrosine | 93829-25G, Sigma Aldrich | 133 |
| L-Valine  Adenine  Cytosine  Uracil  Guanine | 72-18-4, Acros  1152, Calbiochem  71-30-7, Alfa Aesar  6630, Calbiochem  73-40-5, Alfa Aesar | 126  50  50  50  50 |
| **Other Components (g/L)** | | |
| Polysorbate 80 (Tween 80)  Ergosterol | 59924-1KG-F, Fluka  E6510-10G, Sigma | 1  10 |

### Supplemental Table S2. Concentrations of lignocellulose-derived inhibitors in SynHv4.1 medium.

| **Lignocellulose-Derived**  **Inhibitors (LCDIs)** | **Source (Cat. # or CAS #, Vendor)** | | **Concentration in**  **μM** |
| --- | --- | --- | --- |
| **Amides, Imidazoles and Pyrazines (μM)** | | | |
| Feruloyl amide | NA^1^ | | 701 |
| Coumaroyl amide | NA^1^ | | 1104 |
| 2-Methylimidazole  4(5)-Methylimidazole  2, 4-Dimethylimidazole  2-Methylpyrazine  2, 6-Dimethylpyrazine | M50850-100G, Sigma  199885-50G, Sigma  A11949, Alfa Aesar  W330906-100G-K, Sigma  W327301-50G-K, Sigma | | 73  164  7.6  38  14.3 |
| **Acids (μM)** | | | |
| *p*-Coumaric acid | C9008-25G, Sigma | | 146 |
| Ferulic acid | W518301-100G, Sigma | | 33 |
| Benzoic acid | 242381-100G, Sigma | | 344 |
| Vanillic acid | H36001-25G, Sigma | | 134 |
| 4-hydroxybenzoic acid | 240141-50G, Sigma | | 62 |
| **Alcohols, Aldehydes and Ketones (μM)** | | | |
| Vanillin | V1104-100G, Sigma | | 84 |
| 4-Hydroxybenzeldehyde | S7602-25G, Sigma | | 70 |
| Coniferyl alcohol | 223735-1G, Sigma | | 55 |
| 4-Hydroxyacetophenone | 278564-100G, Sigma | | 33 |
| Furfural | F94-500, Fisher | | 47 |
| ^1^ [(Keating et al., 2014)](https://doi.org/10.3389/fmicb.2014.00402) |  |  |  |

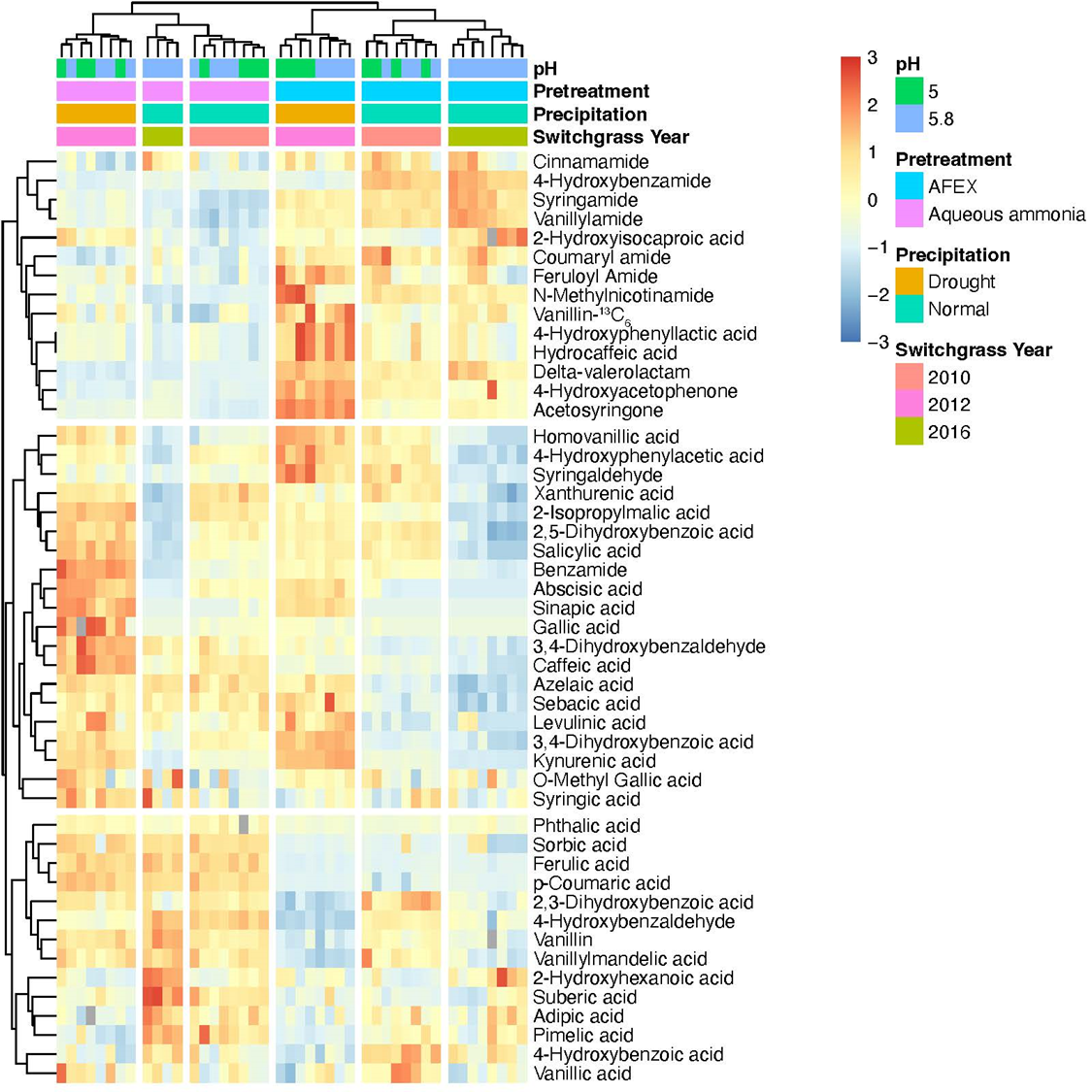
**Fig S1. Heatmap of all 48 compounds quantified by mass spectrometry in AFEX- and SAA-pretreated hydrolysates.** Concentrations are shown as z-scores. Outliers >3 standard deviations from the population mean were omitted from hierarchical clustering and are plotted in grey. Full analyte quantification data can be found in Supplemental File S1.

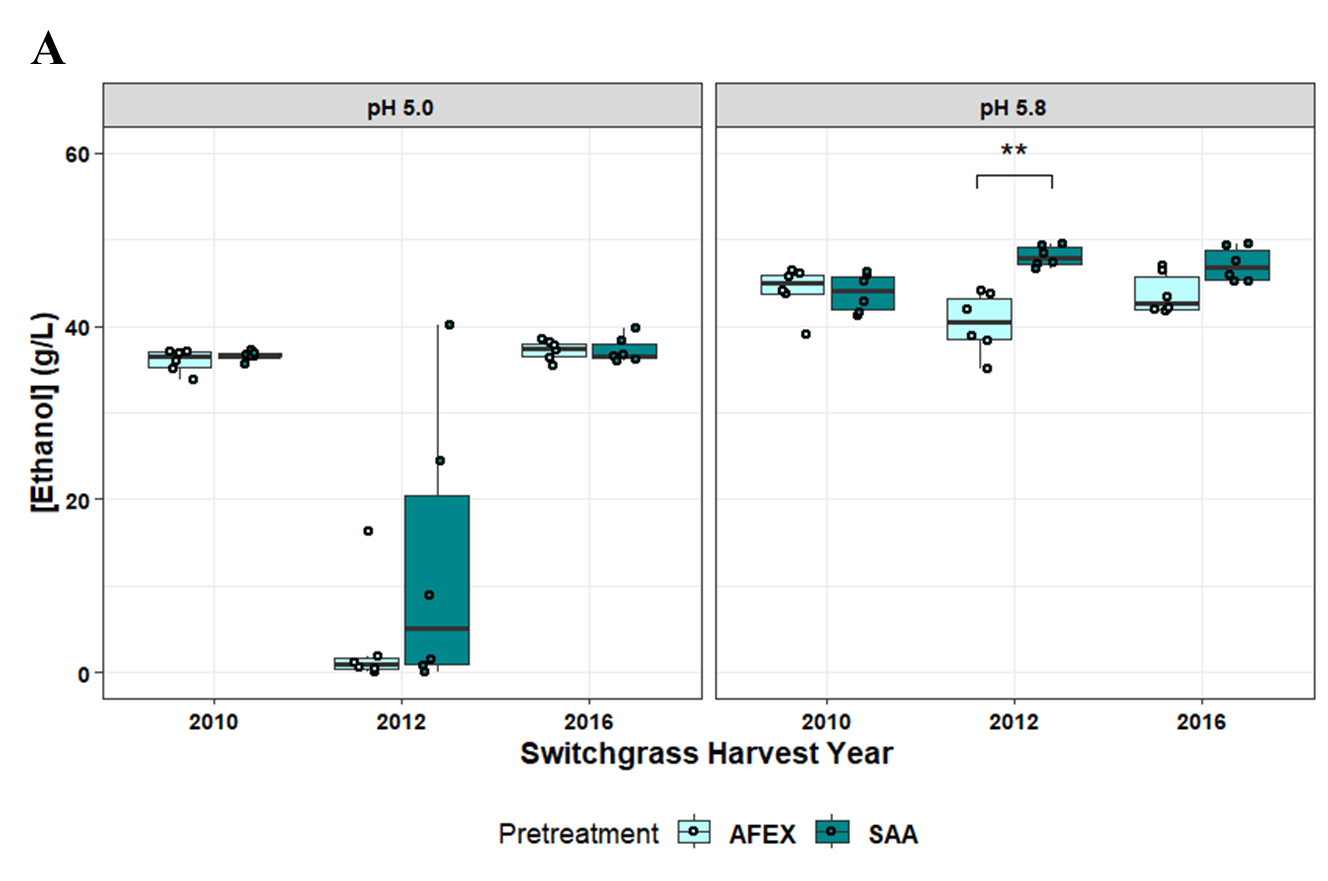

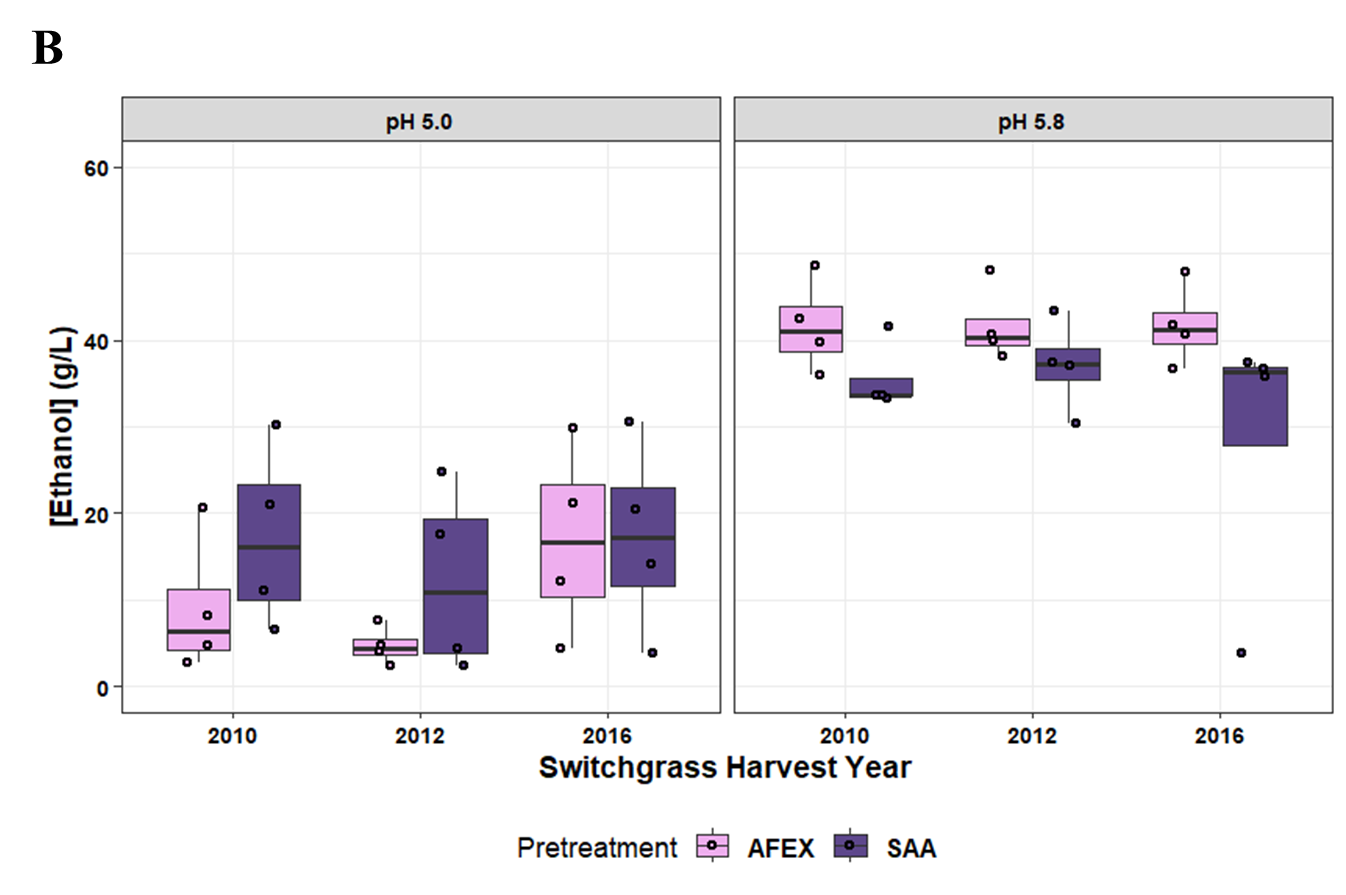
**Figure S2. Final Ethanol titers from Y1455 and Zm2032 fermentations.** Ethanol titers for yeast Y1455 (**A**) and Zm2032 (**B**) from Fig.3 grouped by pH. Brackets denote significance as determined by two-sided Wilcoxon test; * p<0.05, **p<0.01, ***p<0.001.

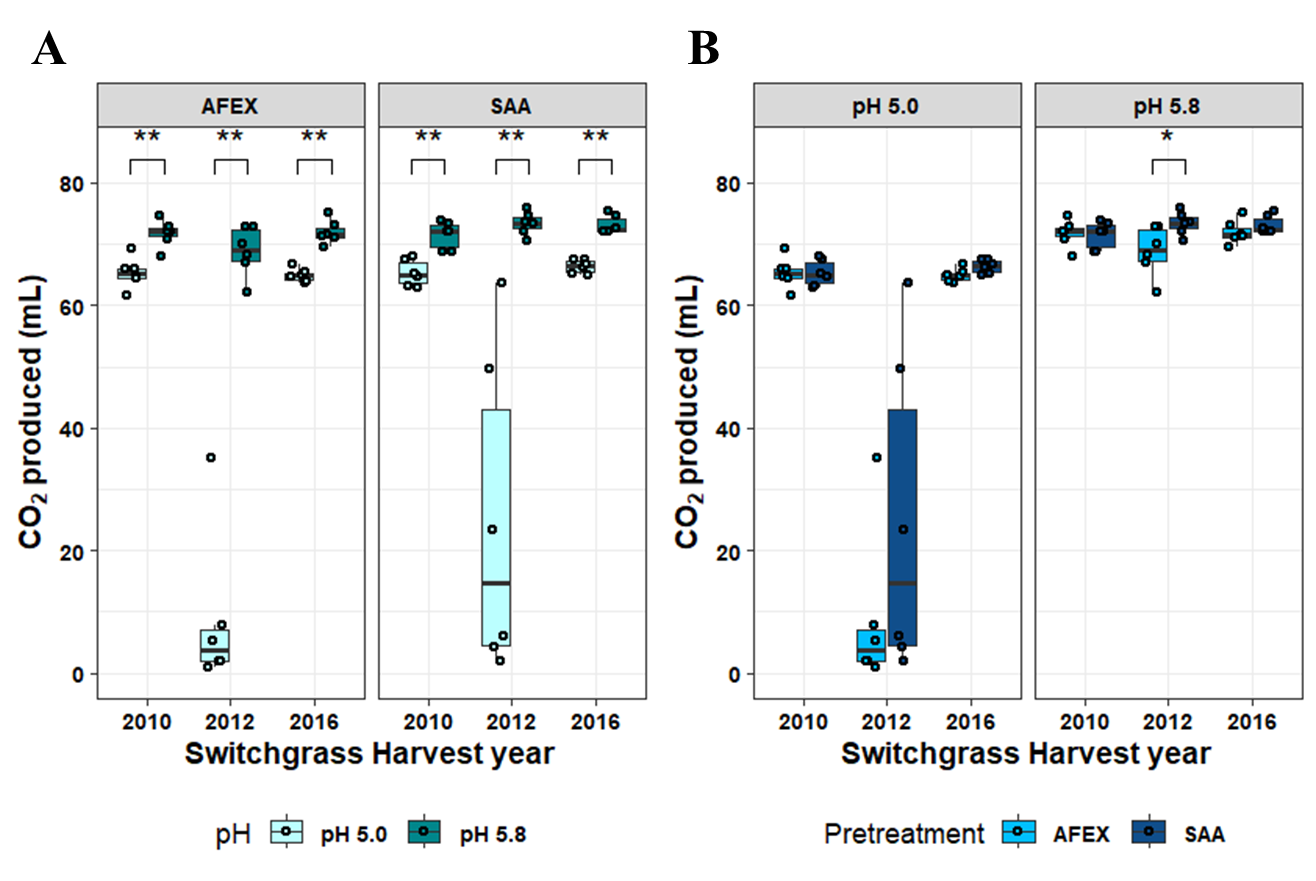

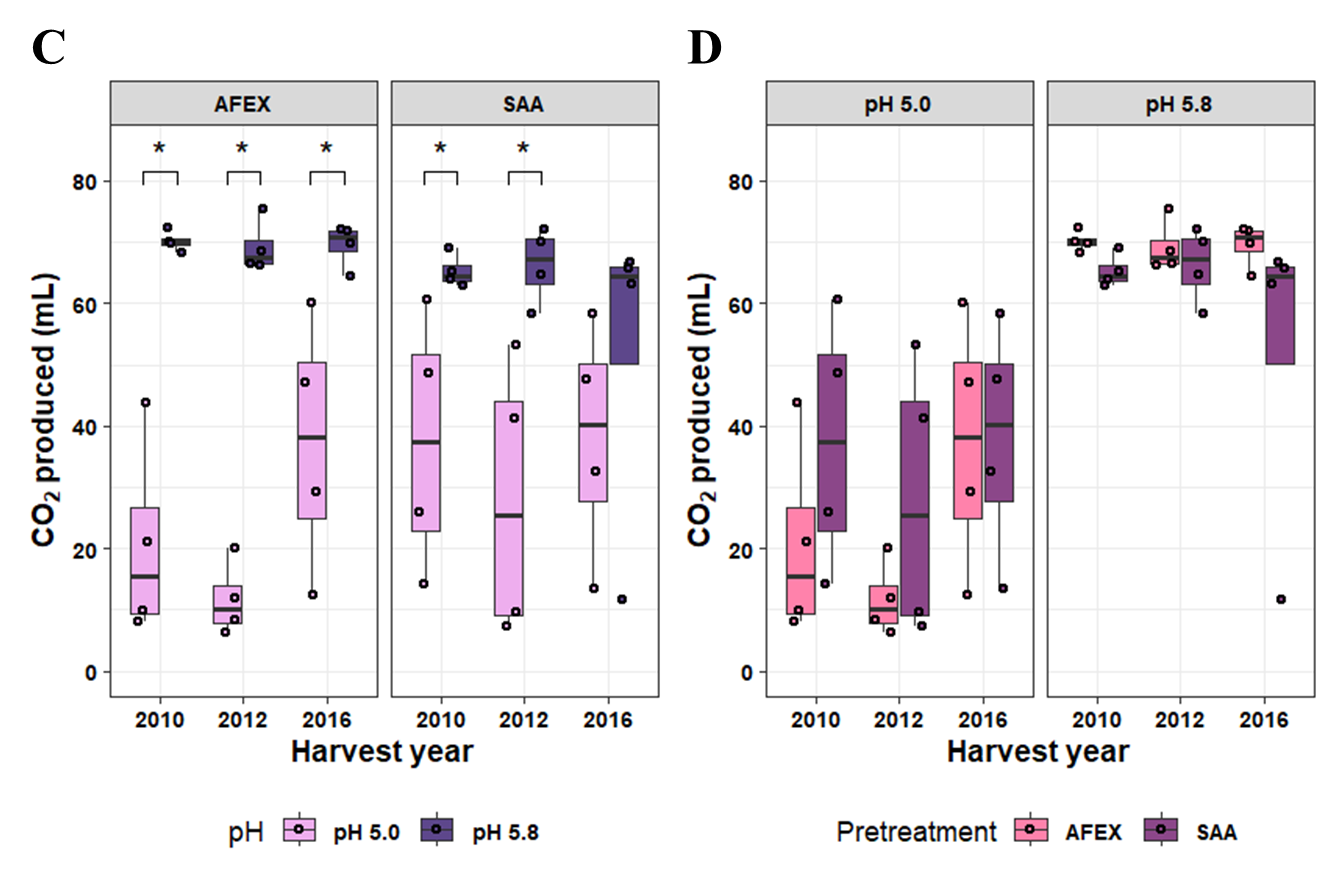
**Figure S3. Boxplots of total CO_2_ produced by yeast and bacteria.** CO_2_ volumes produced by Y1455 **(A, B**) and Zm2023 (**C, D**) are grouped by pretreatment (**A, C**) and by pH (**B, D**) to illustrate relevant comparisons. Brackets denote significance level from two-sided Wilcoxon tests within each switchgrass harvest year; * p<0.05, **p<0.01, ***p<0.001.

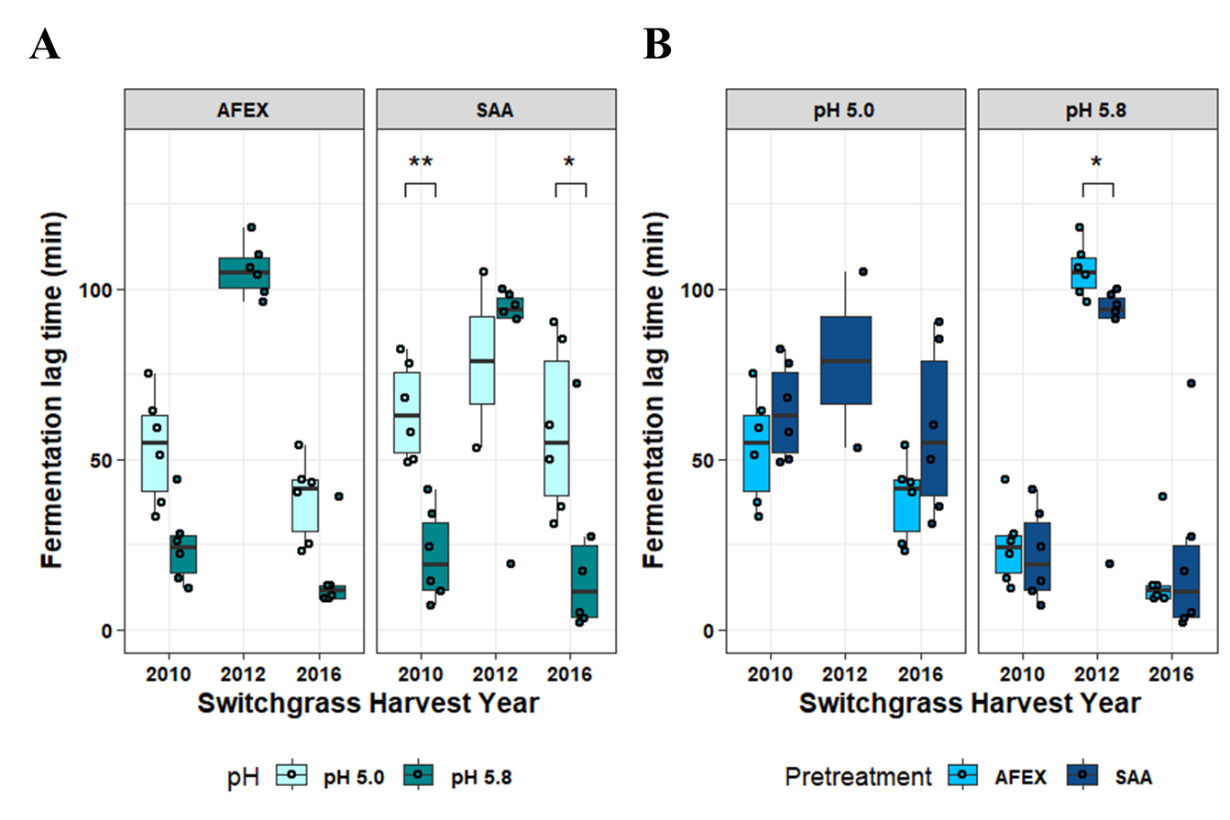

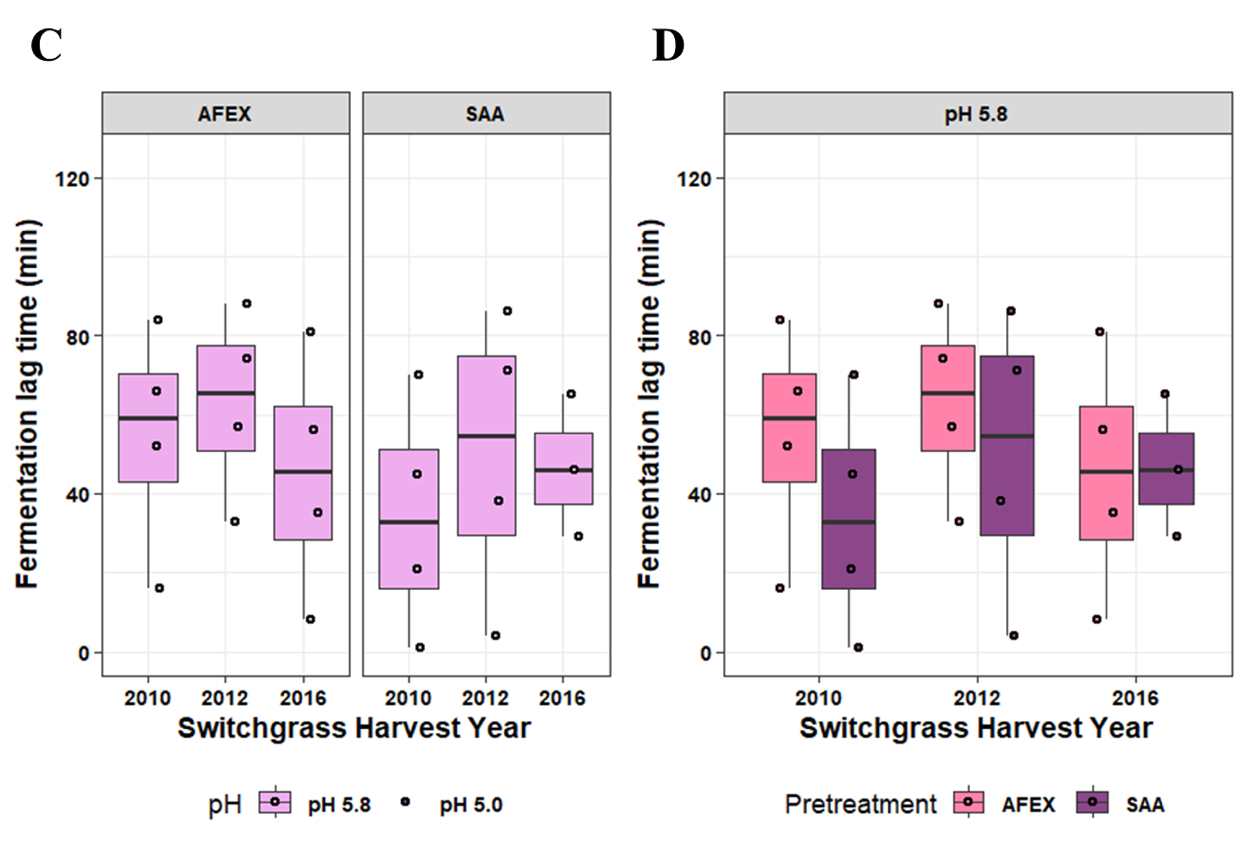

**Figure S4. Lag time for fermentations of AFEX and SAA hydrolysates by** **Y1455 and Zm2032.** For Y1455 (**A, B**) and Zm2032 (**C,D**), fermentation lag time data is plotted grouped by pretreatment (**A, C**) and by pH (**B, D**) to illustrate relevant comparisons. Brackets denote significance level from two-sided Wilcoxon tests within each switchgrass harvest year; * p<0.05, **p<0.01, ***p<0.001. Note that lag time could not be accurately estimated for severely inhibited fermentations, leading to missing data.

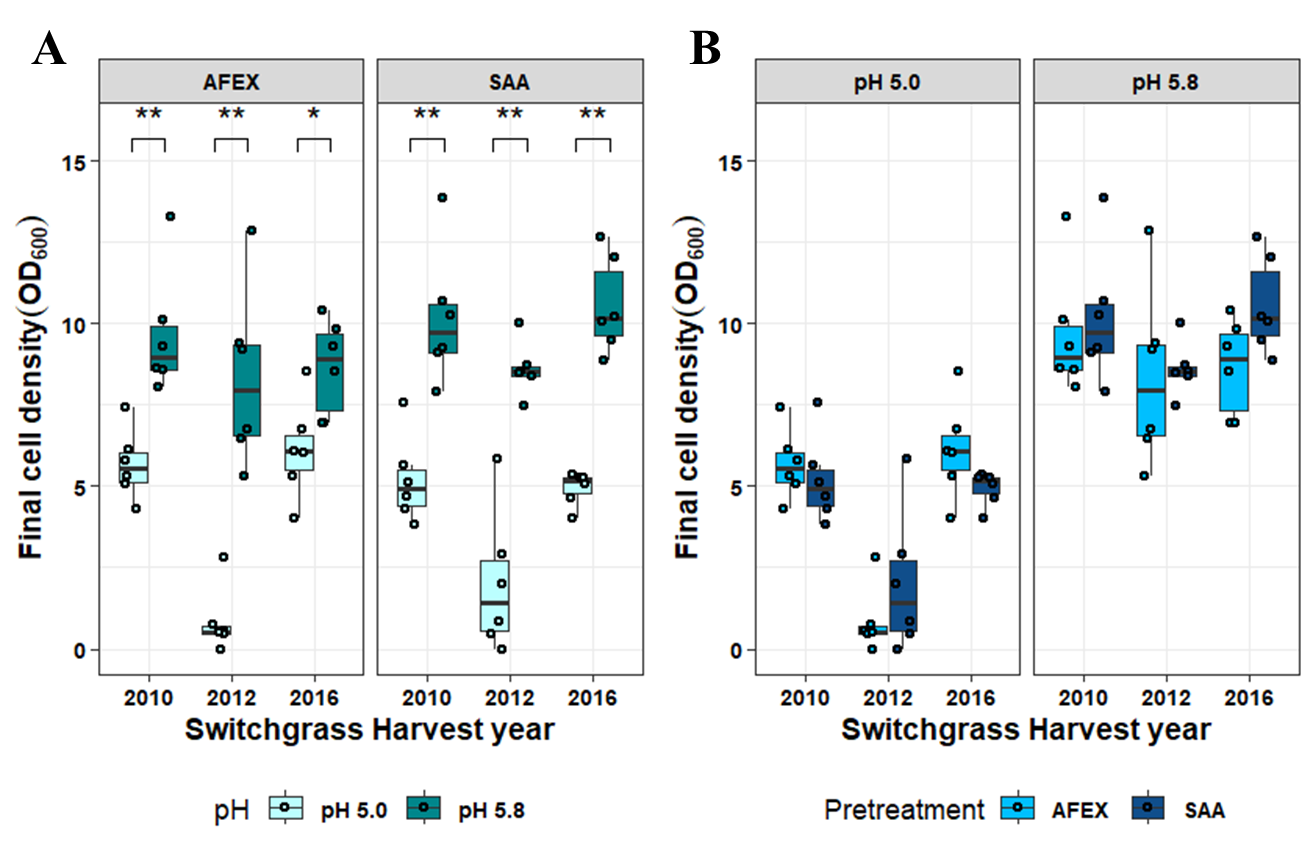

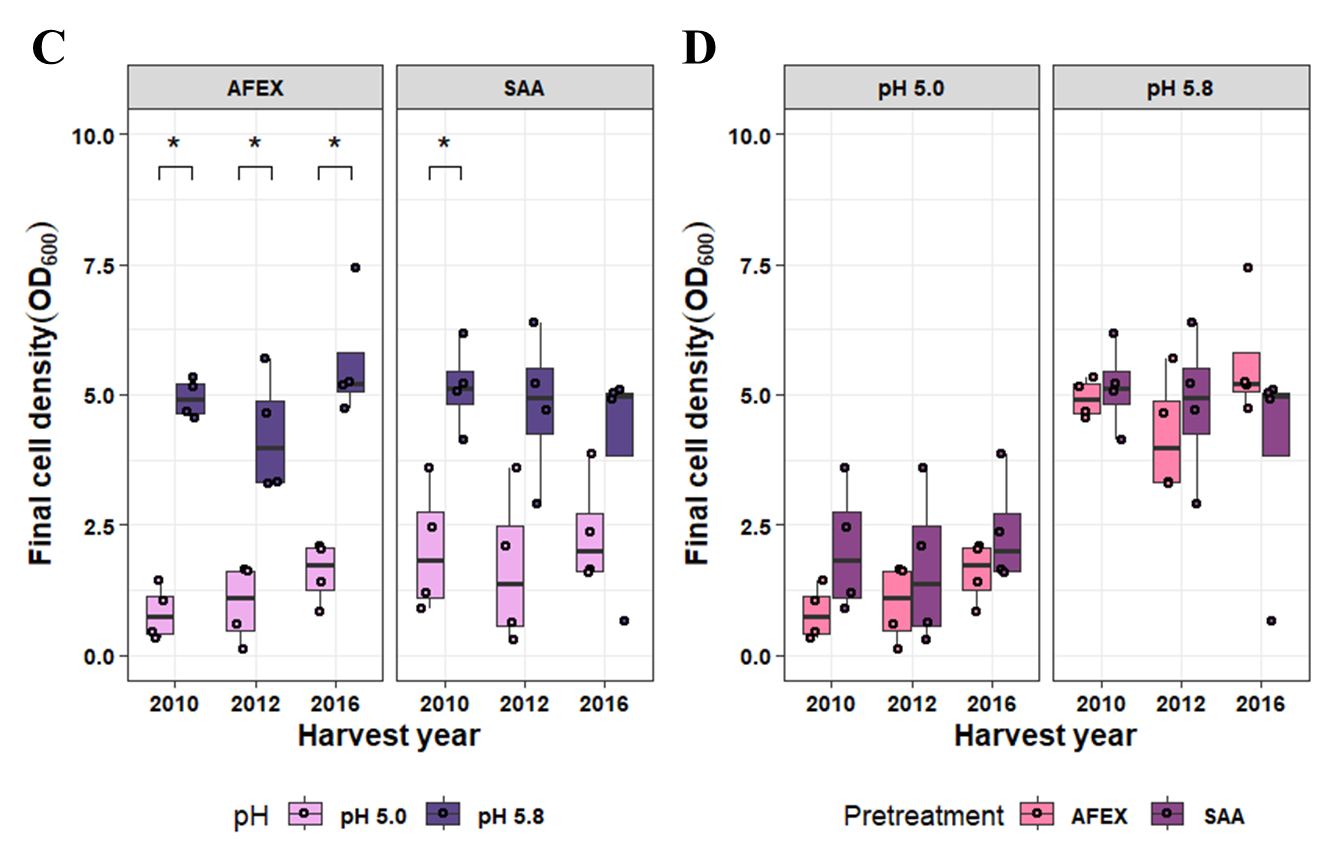
**Figure S5. Boxplots of final cell densities for Y1455 and Zm2032.** For Y1455 (**A, B**) and Zm2032 (**C,D**), final cell densities are grouped by pretreatment (**A, C**) and by pH (**B, D**) to illustrate relevant comparisons. Brackets denote significance level from two-sided Wilcoxon tests within each switchgrass harvest year; * p<0.05, **p<0.01, ***p<0.001

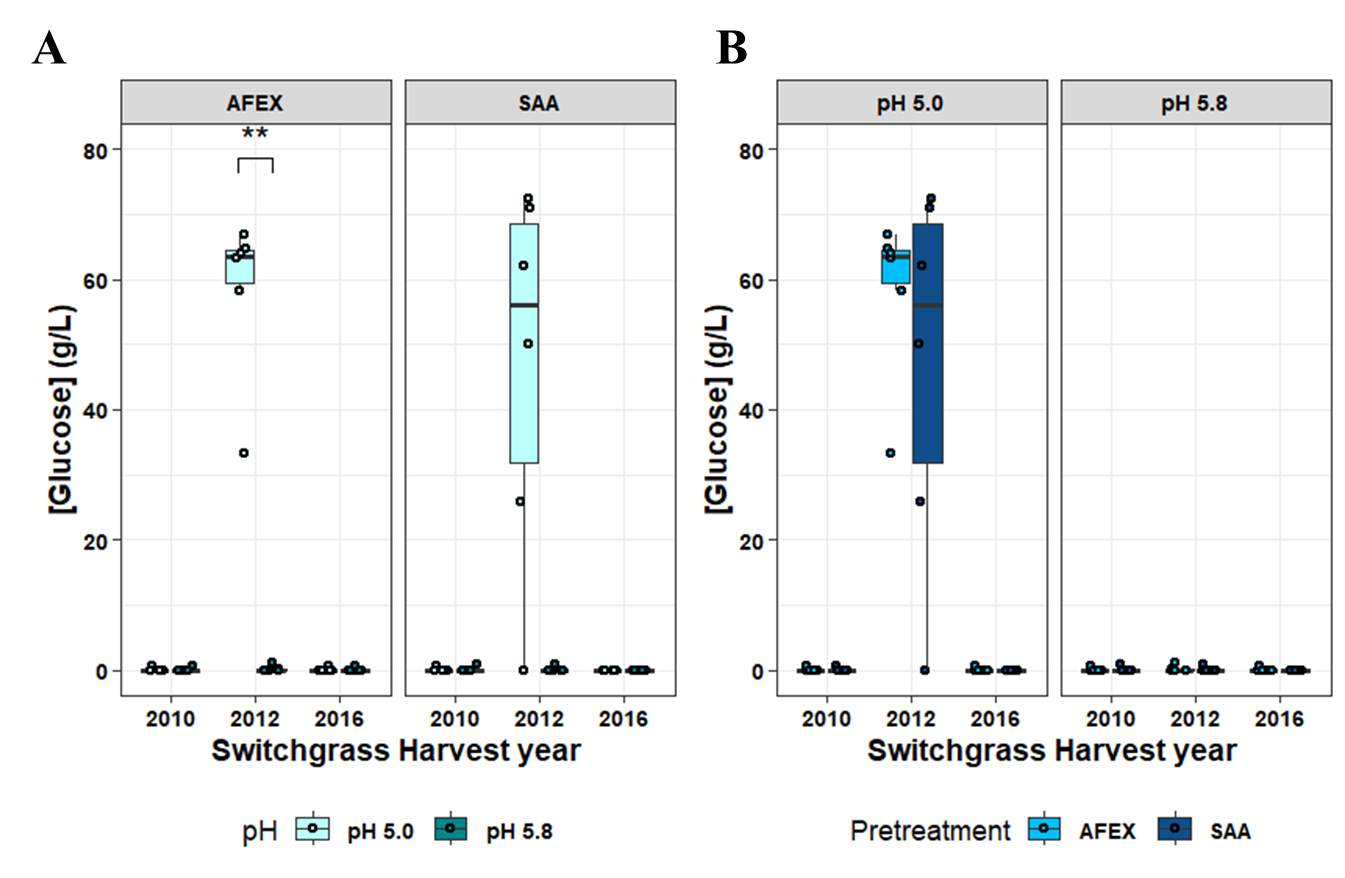

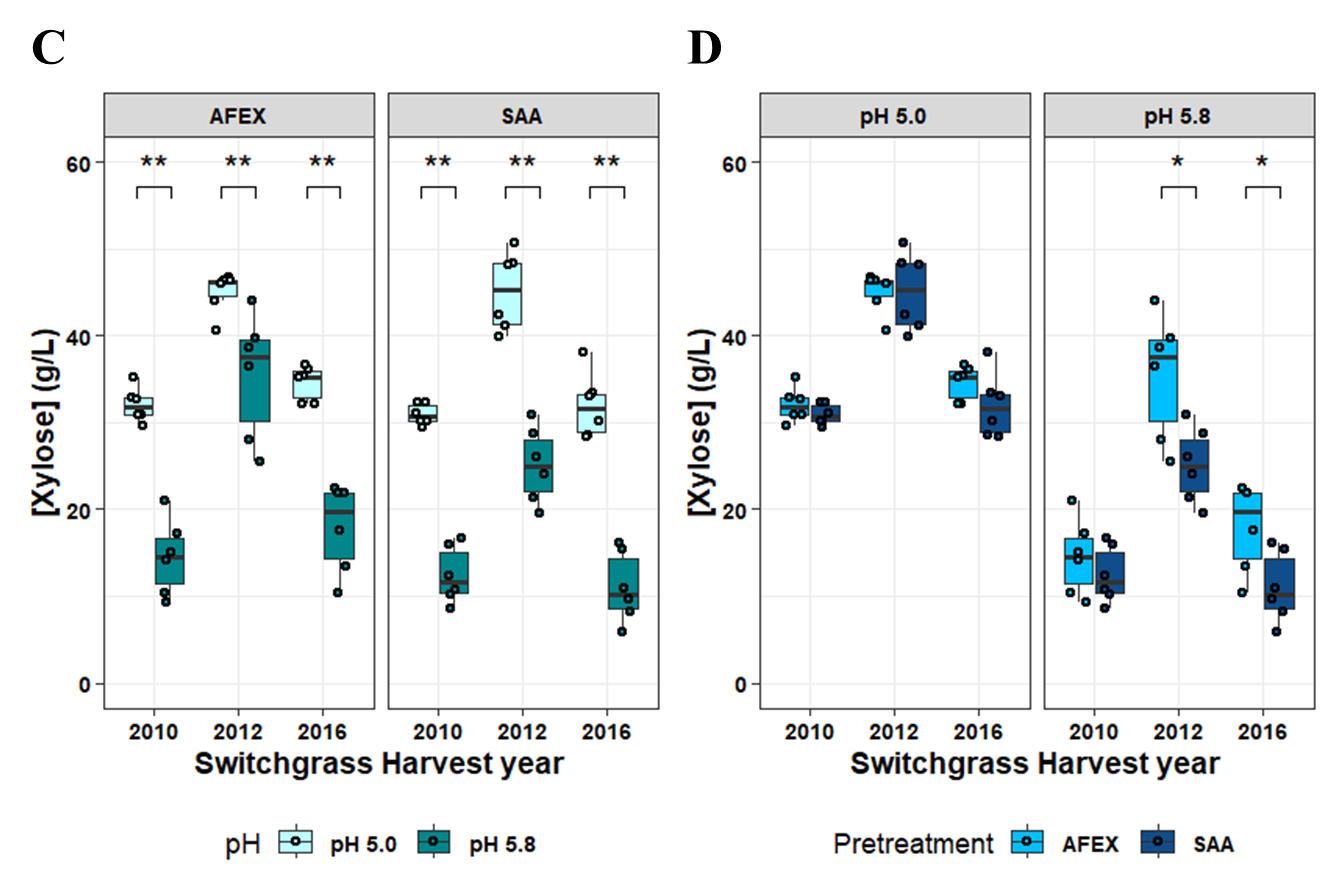

**Figure S6. Boxplots of glucose and xylose remaining after fermentation of hydrolysates by Y1455 yeast.** Final sugar concentrations are grouped by pretreatment (**A, C**) and by pH (**B, D**) to illustrate relevant comparisons. Brackets denote significance level from two-sided Wilcoxon tests within each switchgrass harvest year; * p<0.05, **p<0.01, ***p<0.001.

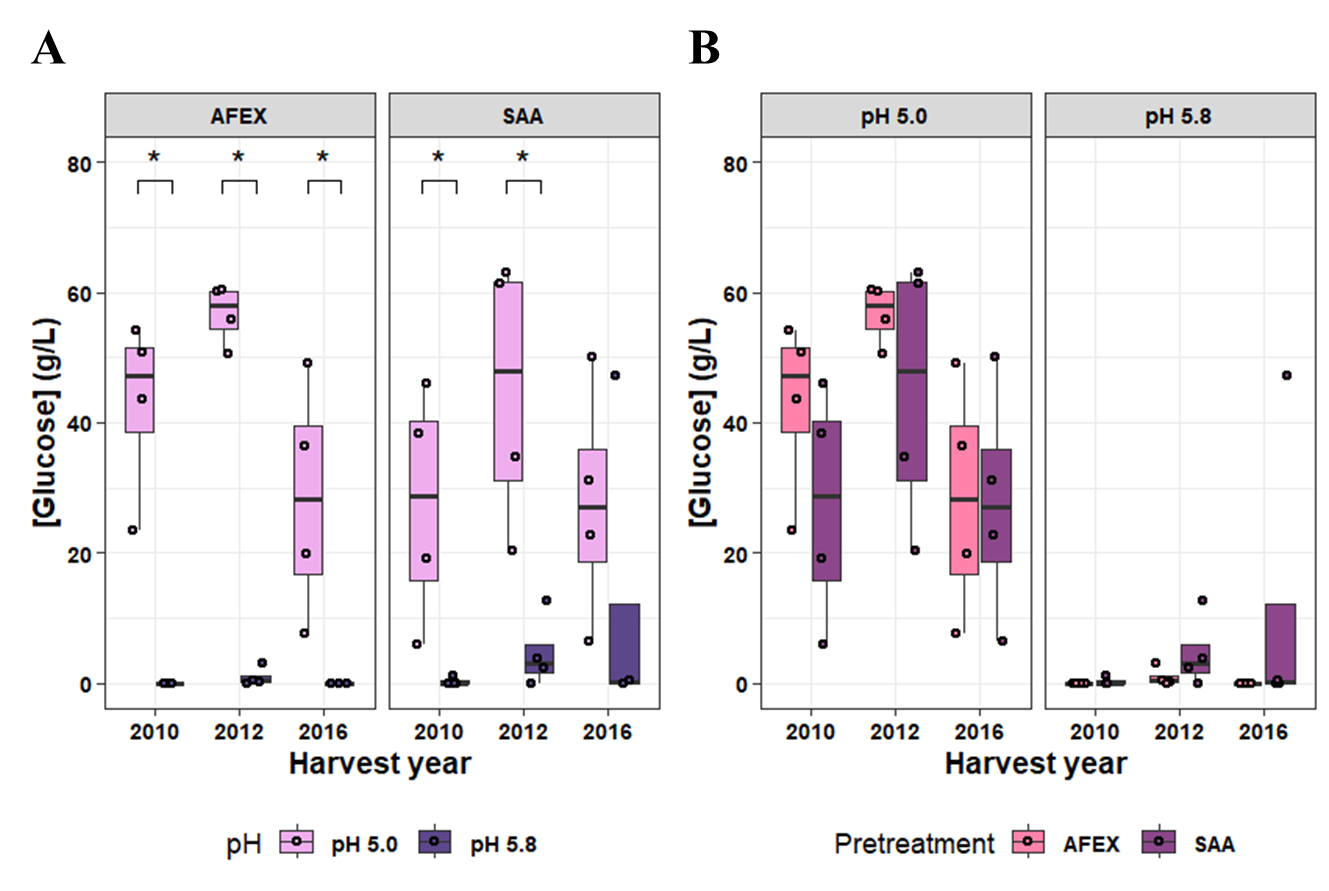

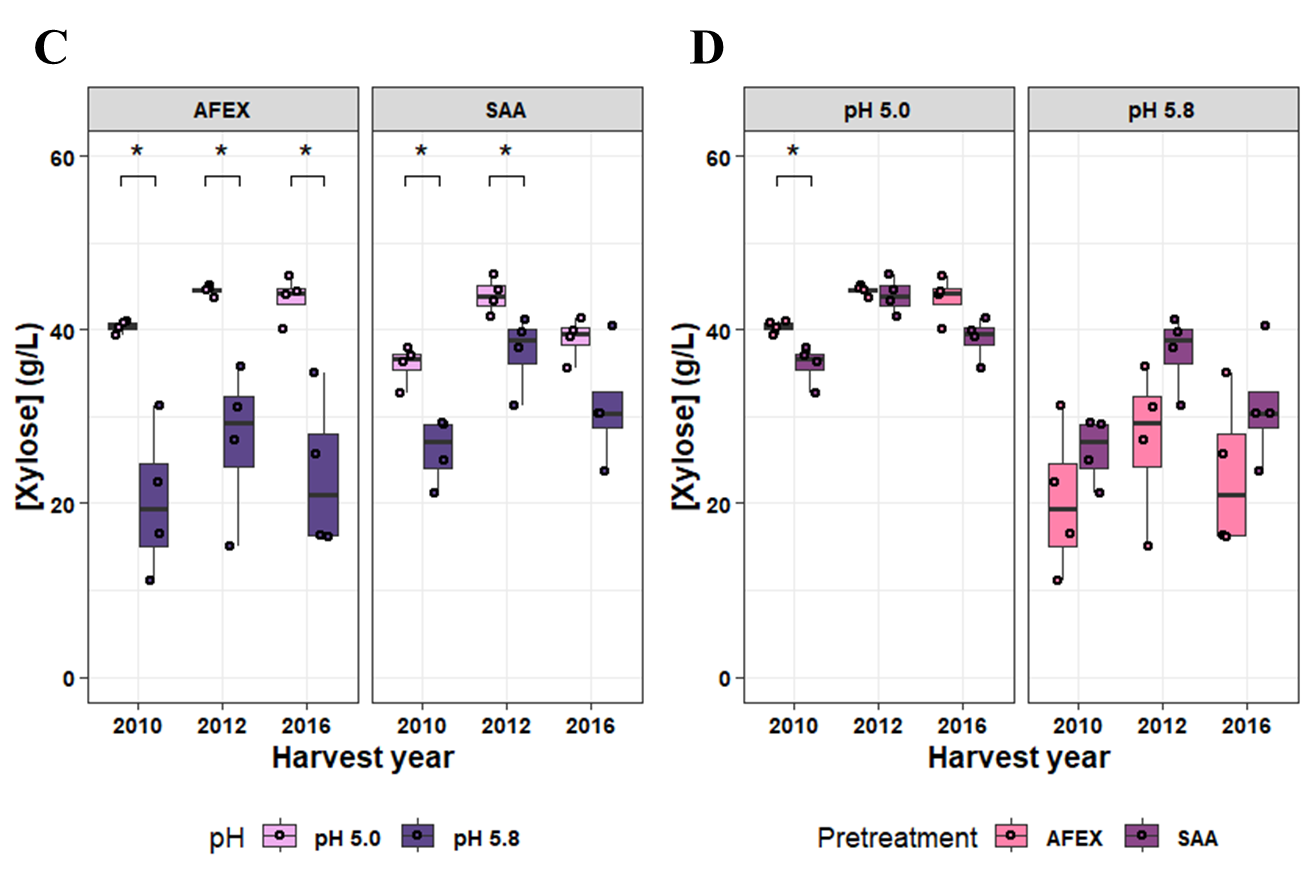

**Figure S7. Boxplots of glucose and xylose remaining after fermentation of hydrolysates by Zm2032 bacteria.** Final sugar concentrations are grouped by pretreatment (**A, C**) and by pH (**B, D**) to illustrate relevant comparisons. Brackets denote significance level from two-sided Wilcoxon tests within each switchgrass harvest year; * p<0.05, **p<0.01, ***p<0.001

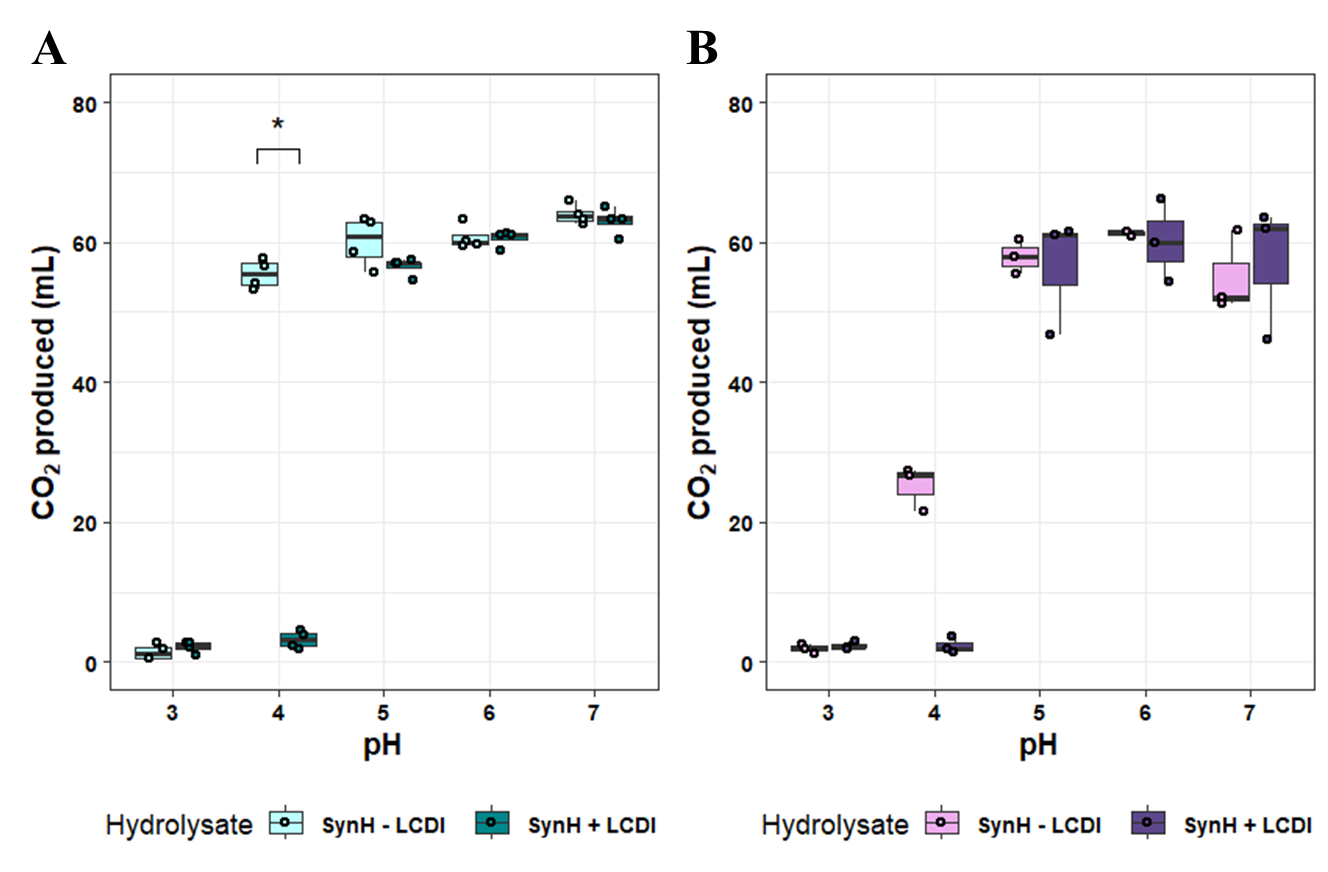
**Figure S8. Boxplots of final CO_2_ produced from fermentations of synthetic hydrolysates by Y1455 and Zm2032.** CO_2_ volumes produced by Y1455 yeast (**A**) and Zm2032 (**B**) fermentations from the indicated synthetic hydrolysates at the indicated pH are plotted. Brackets denote significance level from two-sided Wilcoxon tests between SynH versions at each pH; * p<0.05, **p<0.01, ***p<0.001.

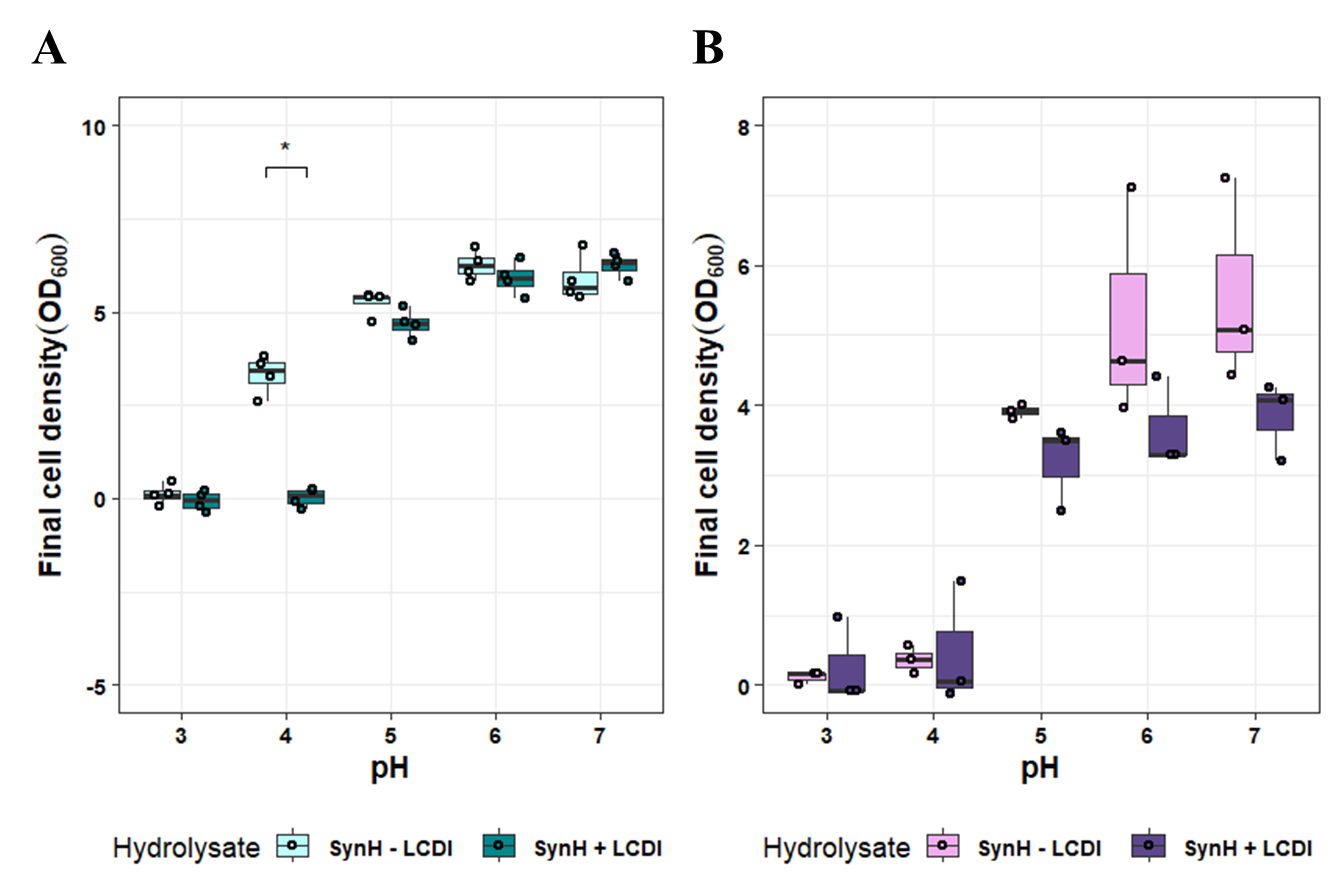
**Figure S9. Boxplots of final cell densities from fermentations of synthetic hydrolysates by Y1455 and Zm2032.** For Y1455 (**A, B**) and Zm2032 (**C,D**), final cell densities are plotted by medium pH. Brackets denote significance level from two-sided Wilcoxon tests between SynH versions at each pH; * p<0.05, **p<0.01, ***p<0.001.

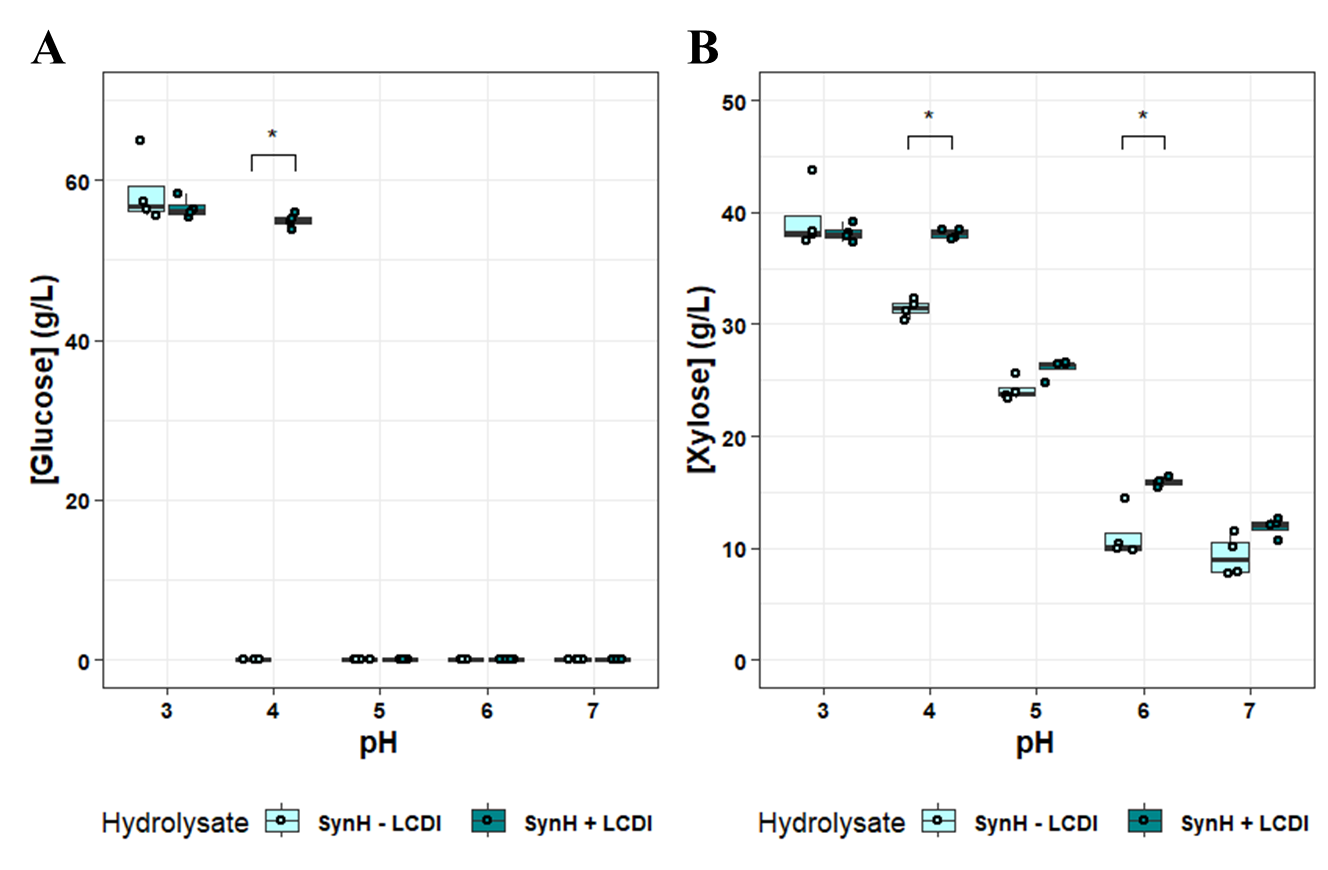

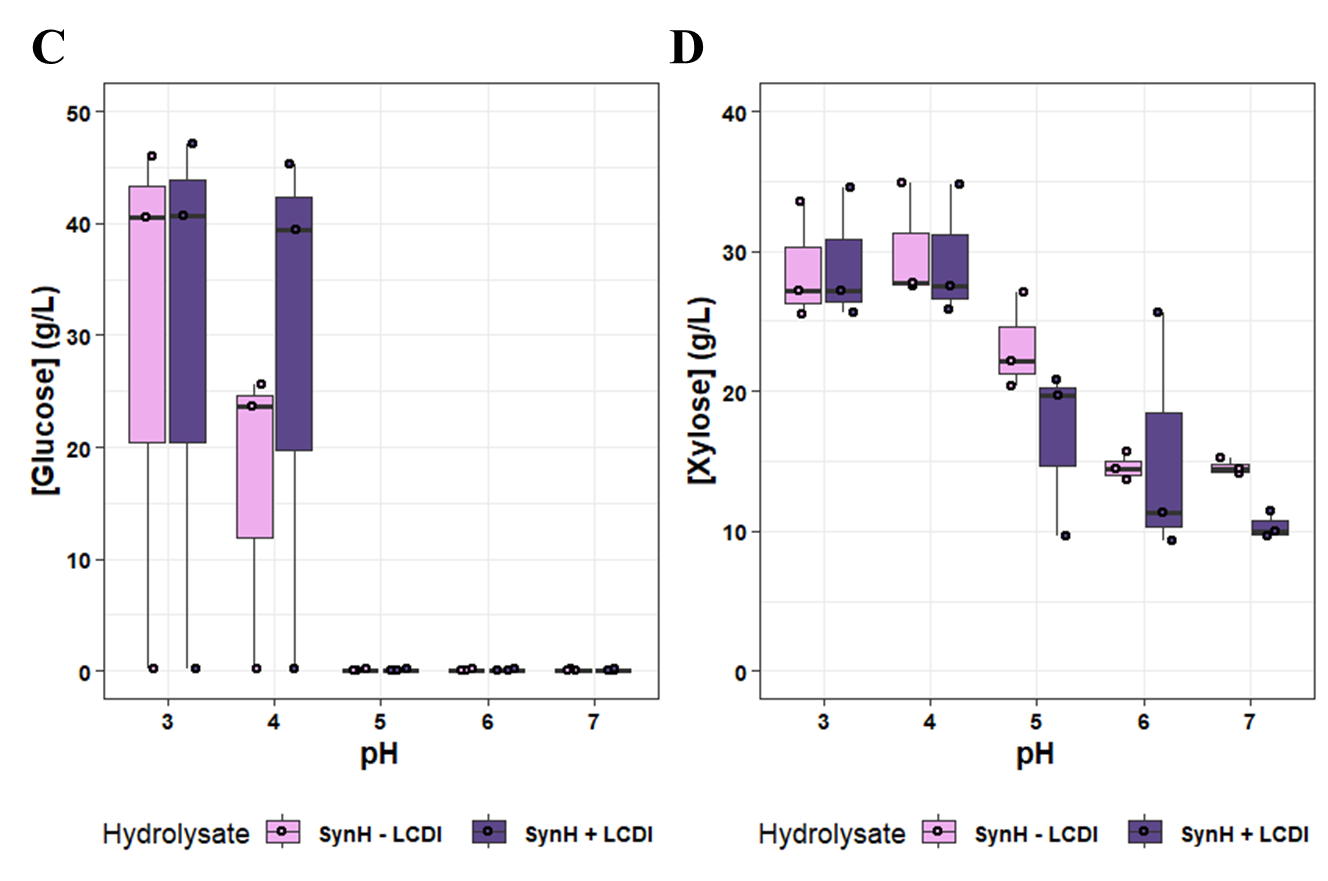

**Figure S10. Boxplots of remaining glucose and xylose following fermentations of synthetic hydrolysates by Y1455 and Zm2032.** Final glucose and xylose concentrations from Y1455 (**A, B**) and Zm2032 (**C,D**) fermentations of synthetic hydrolysates are plotted, Brackets denote significance level from two-sided Wilcoxon tests between SynH versions at each pH; * p<0.05, **p<0.01, ***p<0.001.

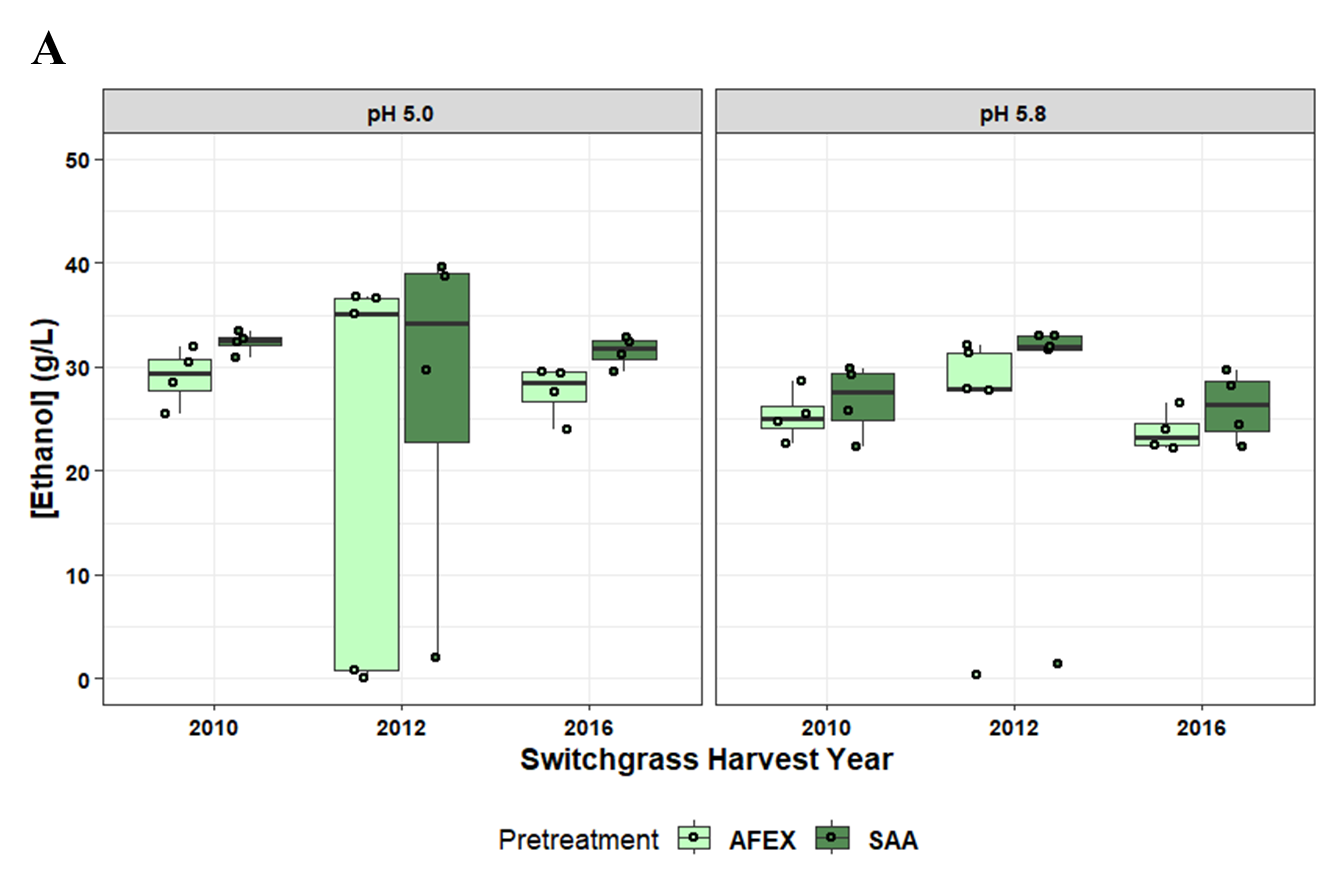

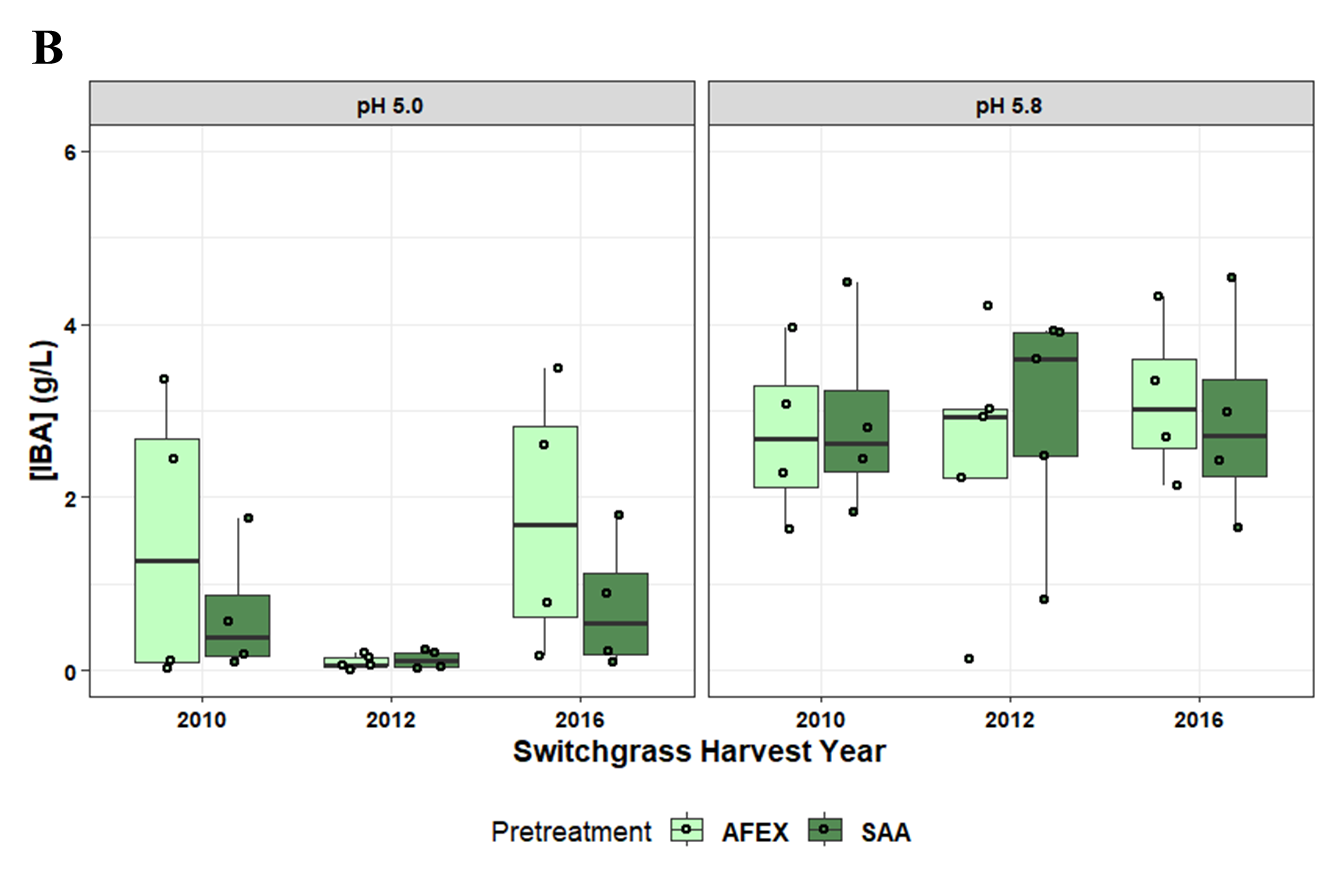
**Figure S11. Final ethanol and isobutanol (IBA) titers by yHRW253.** Ethanol titers using data from Fig. 5 are grouped by pH.

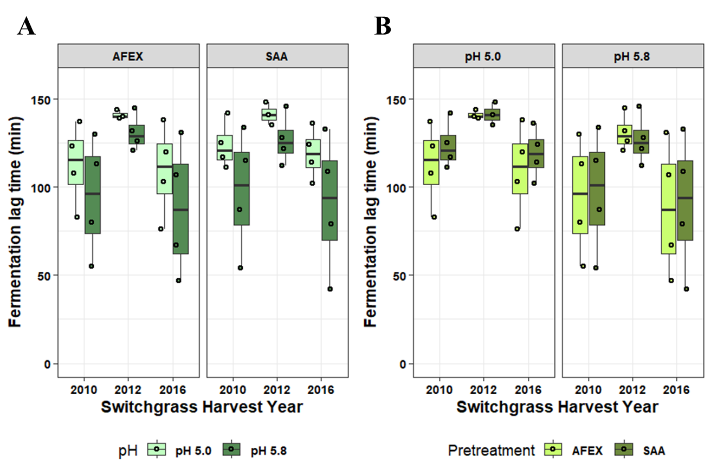

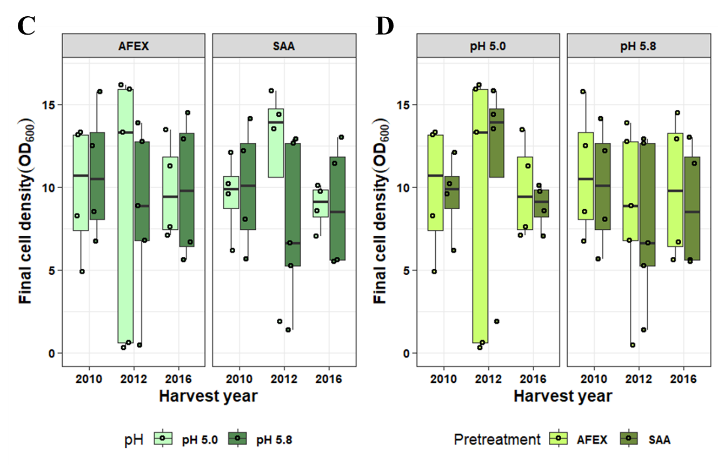

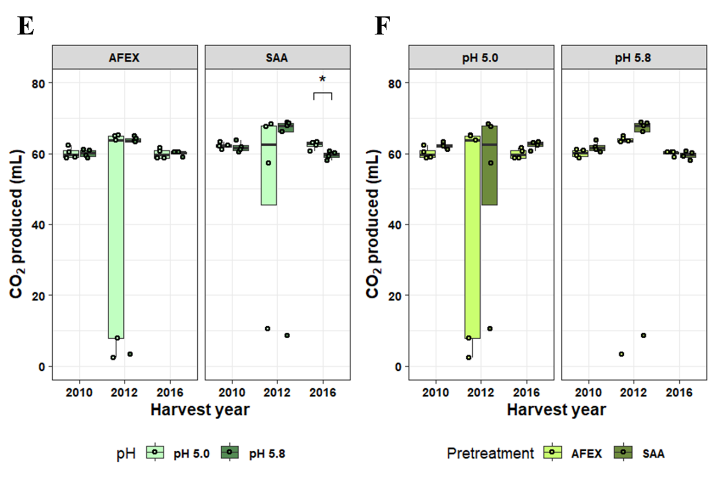

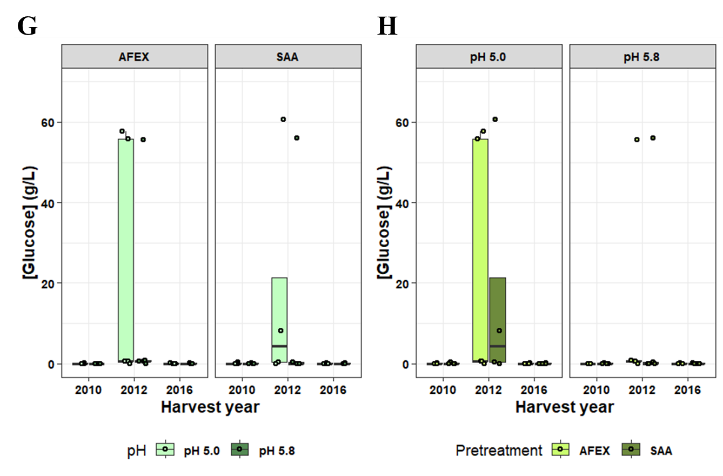

**Figure S12. Boxplots of lag time, final cell densities, CO_2_ produced and glucose concentrations for yHRW253 fermentations.** For each parameter, the same data is plotted grouped by pretreatment (**A, C, E, G**) and by pH (**B, D, F, H**) to illustrate relevant comparisons. Brackets denote significance level from two-sided Wilcoxon tests within each switchgrass harvest year; * p<0.05, **p<0.01, ***p<0.001.
